# Chronic sodium bromide treatment relieves autistic-like behavioral deficits in three mouse models of autism

**DOI:** 10.1101/2021.09.14.460257

**Authors:** Cécile Derieux, Audrey Léauté, Agathe Brugoux, Déborah Jacaz, Jean-Philippe Pin, Julie Kniazeff, Julie Le Merrer, Jerome AJ Becker

## Abstract

Autism Spectrum Disorders (ASD) are neurodevelopmental disorders whose diagnosis relies on deficient social interaction and communication together with repetitive behavior. To date, no pharmacological treatment has been approved that ameliorates social behavior in patients with ASD. Based on the excitation/inhibition imbalance theory of autism, we hypothesized that bromide ions, long used as an antiepileptic medication, could relieve core symptoms of ASD. We evaluated the effects of chronic sodium bromide (NaBr) administration on autistic-like symptoms in three genetic mouse models of autism: *Oprm1^-/-^* , *Fmr1^-/-^* and *Shank3^Δex13-16-/-^* mice. We showed that chronic NaBr treatment relieved autistic-like behaviors in these three models. In *Oprm1^-/-^* mice, these beneficial effects were superior to those of chronic bumetanide administration. At transcriptional level, chronic NaBr in *Oprm1* null mice was associated with increased expression of genes coding for chloride ions transporters, GABAA receptor subunits, oxytocin and mGlu4 receptor. Lastly, we uncovered synergistic alleviating effects of chronic NaBr and a positive allosteric modulator (PAM) of mGlu4 receptor on autistic-like behavior in *Oprm1^-/-^* mice. We evidenced in heterologous cells that bromide ions behave as PAMs of mGlu4, providing a molecular mechanism for such synergy. Our data reveal the therapeutic potential of bromide ions, alone or in combination with a PAM of mGlu4 receptor, for the treatment of ASDs.

## INTRODUCTION

Autism Spectrum Disorders (ASD) are neurodevelopmental diseases with high heterogeneity and heritability. Their diagnostic is reached in presence of impaired social communication and interaction together with a restricted, repetitive repertoire of behaviors, interests and activities (*1*). Alongside core symptoms, ASD are often associated with neurobehavioral comorbidities, such as high anxiety, cognitive and motor deficits or epilepsy (*2–5*). Despite the identification of vulnerability genes and environmental risk factors (*6–8*), the etiology of ASD remains essentially unknown, making the development of pharmacological treatments for these pathologies a true challenge.

Excitation/inhibition (E/I) imbalance appears as a common mechanistic feature in ASD (*9, 10*). The heuristic hypothesis of excessive E/I ratio in ASD was initially formulated by Rubenstein and Merzenich (*11*) and raised significant interest as accounting well for reduced GABAergic signaling (*12, 13*) and high prevalence of epilepsy (10-30%) (*3*) in these pathologies. Indeed, epilepsy is one of the most frequent comorbid medical condition in autism (*5, 14*) and the prevalence of epileptiform EEG or altered resting-state is even higher (*15, 16*), suggesting shared risk factors and/or pathophysiological mechanisms (*17, 18*). However, the excessive E/I hypothesis in ASD has been challenged by studies in animal models showing instead decreased excitation, which led to a more general concept of altered E/I homeostasis (*10, 19*).

Compromised E/I balance in ASD may result from several neuropathological mechanisms. On the excitation side, glutamatergic transmission was found altered both in patients and animal models, although in different directions depending on genetic mutations/models (*9, 20, 21*). On the inhibition side, and consistent with impaired GABAergic signaling, decreased levels of GABA (*22*) and expression of GABAA and GABAB (*23, 24*) receptors as well as genetic polymorphisms in GABAA receptor subunits (*25, 26*) have been detected in patients with autism. Accordingly, decreased GABAergic neurotransmission has been reported in several ASD models (*27–31*). Moreover, preclinical studies showed that low doses of benzodiazepines, behaving as positive allosteric modulators (PAMs) of the GABAA receptor (*31, 32*), or the GABAB receptor agonist arbaclofen improve autistic-like behaviors in animal models (*33, 34*). Disappointingly, these results failed to translate to Fragile X syndrome in clinical trials (*35, 36*). Alternatively, it was proposed that GABA neurons remain immature in ASD, failing to shift from high to low intracellular concentrations of chloride ion (Cl^-^), resulting in maintained depolarizing Cl^-^ efflux through activated GABAA receptor (*37*). Intracellular Cl^-^ concentration is under the control of the main Cl^-^ importer NKCC1 (Na^+^-K^+^-2Cl^-^ cotransporter) and the main chloride exporter KCC2. Therefore blocking NKCC1 using the loop diuretic and antiepileptic drug (*38, 39*) bumetanide appeared a promising therapeutic approach in ASD. Accordingly, bumetanide improved autistic-like phenotype in rodent models of ASD (*40*) and relieved autistic behavior in small cohorts of patients (*41, 42*) but failed to demonstrate clinical benefit in a larger clinical trial, except for a reduction of repetitive behavior (*43*).

Bromide ion (Br^-^) was the first effective treatment identified for epilepsy (*44*), long used also as an anxiolytic and hypnotic medication (*45*). With the advent of novel antiepileptic and anxiolytic drugs, the use of Br^-^ was progressively dropped down, although it remains a valuable tool to treat refractory seizures (*46, 47*). As regards its mechanism of action, Br^-^ shares similar chemical and physical properties with Cl^-^, allowing it substituting Cl^-^ in multiple cellular mechanisms. These include anion efflux through activated GABAA receptor, with higher permeability to Br^-^ compared to Cl^-^ resulting in neuronal hyperpolarization (*48*), and transport through the NKCC and KCC cotransporters (*49, 50*). In view of the E/I imbalance theory, these properties point to Br^-^ as an interesting candidate for ASD treatment.

In the present study, we assessed the effects of chronic sodium bromide administration on core autistic-like symptoms: social deficit and stereotypies, as well as on a frequent comorbid symptom: anxiety, in three genetic mouse models of autism: *Oprm1^-/-^* , *Fmr1^-/-^* and *Shank3^Δex13-16-/-^* mice, by means of thorough behavioral assessment. Altered E/I balance and/or modified expression of genes involved in this balance have been reported for these three models (*30, 51–55*); the *Oprm1* knockout model presents the advantage of limited impact on learning performance (*52*), allowing better disentangling autistic features from cognitive deficit. We evidenced that Br^-^ treatment alleviates most of the behavioral deficits observed in these mice, and increases expression of various genes within the social brain circuit. We unraveled that Br- not only increases mGlu4 receptor gene expression but also potentiates the effects of the mGlu4 PAM VU0155041 as well as its agonist glutamate, in *Oprm1^-/-^* mice and in heterologous cells. Our data reveal the therapeutic potential of Br^-^ administration and its combination with a positive allosteric modulator (PAM) of mGlu4 receptor for the treatment of ASD.

## RESULTS

### Chronic sodium bromide was more efficient than bumetanide to relieve social behavior deficits in *Oprm1^-/-^* mice

We first assessed the effects of NaBr administration over a wide range of doses (10 to 500 mg/kg) in *Oprm1^-/-^* mice and their WT counterparts, and compared with bumetanide administration (0.5 and 2 mg/kg) (Fig. 1A). Treatment was given chronically to mimic clinical conditions.

**Fig 1.**
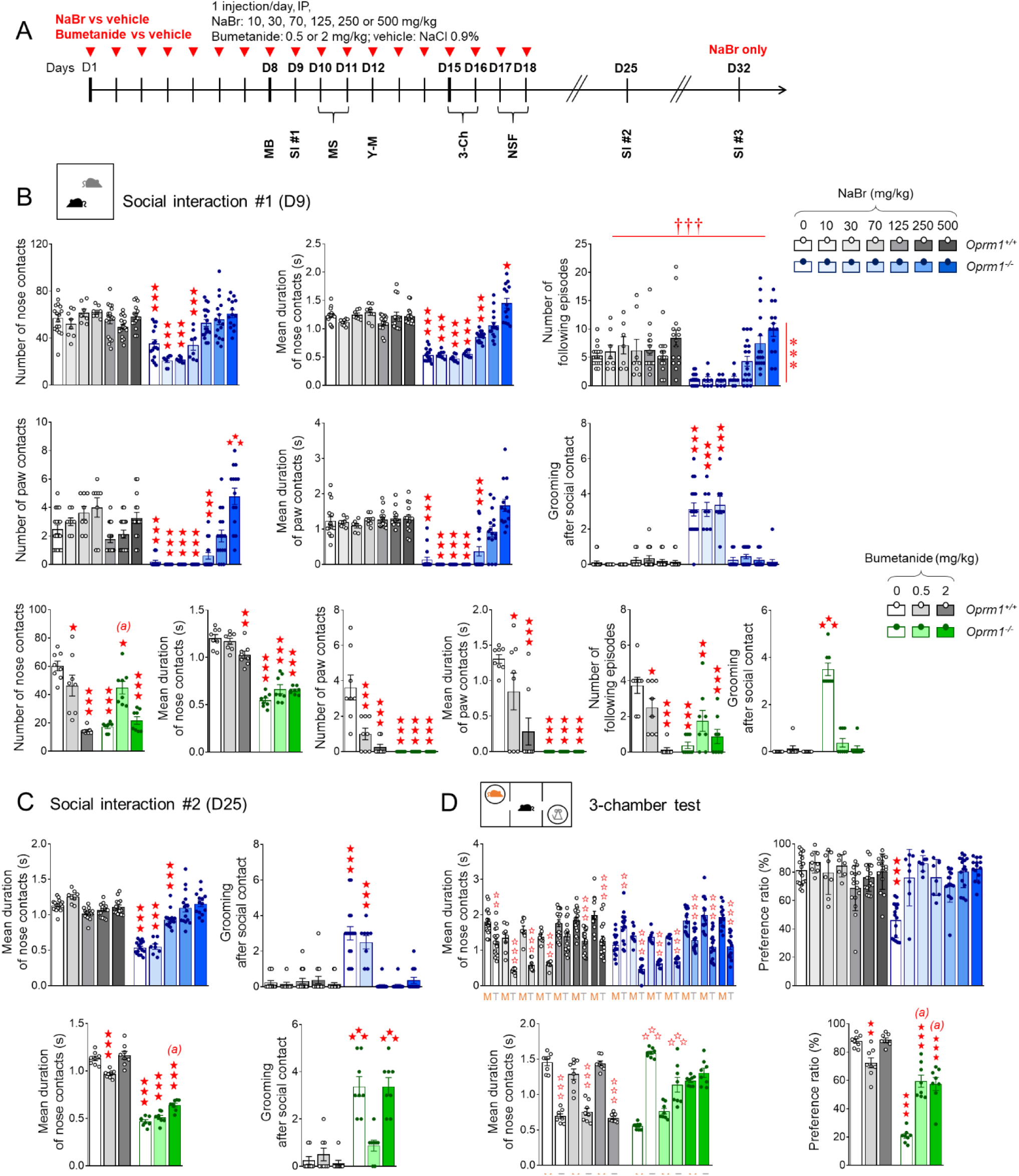
Chronic sodium bromide dose-dependently relieved social behavior deficits in *Oprm1^-/-^* mice, demonstrating superior effects to chronic bumetanide. (A) *Oprm1^+/+^* and *Oprm1^-/-^* mice were treated either with NaBr (0, 125-500 mg/kg: n=14-20 mice per genotype and dose; 10-70 mg/kg: n=8 mice per genotype and dose) or with bumetanide (0, 0.5 and 2 mg/kg: n=8-10 mice per genotype and dose) once daily for 18 days. Behavioral testing started on D8; social interaction was retested 1 week and 2 weeks after cessation of chronic administration. (B) In the direct social interaction test (D9), chronic NaBr administration relieved social deficits of *Oprm1* null mice in a dose-dependent manner for doses over 125 mg/kg; it had no detectable effect in *Oprm1^+/+^* mice. Bumetanide had only partial effects, increasing the number of nose contacts (low dose) and suppressing grooming after social contact; at the highest dose, it impaired social interaction in wild-type controls. (C) One week after cessation of treatment, beneficial effects of bromide administration were preserved for doses over 125 mg/kg; effects of bumetanide one duration of nose contacts and grooming after social contact were still detectable. (D) In the three-chamber test, NaBr treatment rescued social preference in *Oprm1* mutants since the dose of 10 mg/kg, while bumetanide increased their interest for the mouse without reducing their abnormal interest for the object. Results are shown as scatter plots and mean ± sem. Daggers: genotype effect, asterisks: treatment effect, solid stars: genotype x treatment interaction (comparison to wild-type vehicle condition), open stars: genotype x treatment x stimulus interaction (mouse versus object comparison), (a) genotype x treatment interaction (comparison with knockout vehicle condition, p<0.001) (two-way ANOVA or three-way ANOVA with stimulus as repeated measure, followed by Newman-Keuls post-hoc test). One symbol: p<0.05, two symbols: p<0.01; three symbols: p<0.001. More behavioral parameters in Fig. S2. 3-Ch: 3-chamber test, M: mouse, MB: marble burying, MS: motor stereotypies, NSF: novelty-suppressed feeding, SI: social interaction, T: toy, Y-M: Y-maze.

Social interaction was evaluated after 9 days of chronic NaBr treatment (Fig. 1B, more parameters in Fig. S2A). *Oprm1^-/-^* mice exhibited a severe decrease in social interaction; chronic NaBr administration from the dose of 125 mg/kg dose-dependently relieved this deficit in mutant mice, as evidenced by restored number *(genotype x treatment: F_6,163_=13.2, p<0.0001)* and mean duration *(genotype x treatment: F_6,163_=31.6, p<0.0001)* of nose contacts and normalized number *(genotype x treatment: F_6,163_=14.7, p<0.0001)* and mean duration (*genotype x treatment: F_6,163_=14.5, p<0.0001)* of paw contacts. Chronic NaBr also increased the number of following episodes in both mouse lines *(treatment: F_6,163_=5.6, p<0.0001)* and normalized the frequency of grooming after social contact in mutants, since the dose of 70 mg/kg (*genotype x treatment*: *F_6,163_*=32.2, *p<0.0001*). In contrast, a single acute injection of NaBr (250 mg/kg) had little effect on social interaction parameters (Fig. S2B). When given chronically, NaBr (over 70 mg/kg) produced relieving effects that were still detectable one week after cessation of treatment, as evidenced by preserved restoration of the duration of nose contacts (*genotype x treatment*: *F_6,134_*=53.3, *p<0.0001*) and maintained suppression of grooming after social contact (*genotype x treatment*: *F_6,134_*=30.4, *p<0.0001*) for the highest doses (Fig. 1C and S3). These effects had mostly vanished after two weeks (Fig. S3).

Compared with chronic NaBr, chronic bumetanide increased the number of nose contacts *(genotype x treatment: F_2,42_=22.6, p<0.0001)* and following episodes at low dose *(genotype x treatment: F_2,42_=12.9, p<0.0001)* but failed to increase significantly the duration of nose contacts *(genotype x treatment: F_2,42_=22.6, p<0.0001)* or the number *(genotype x treatment: F_2,42_=14.9, p<0.0001)* and duration *(genotype x treatment: F_2,42_=7.3, p<0.0001)* of paw contacts. Finally, bumetanide suppressed grooming episodes, notably those occurring after social contact, since the lowest dose tested *(genotype x treatment: F_2,42_=80.7, p<0.0001)*. Of note, chronic bumetanide treatment showed deleterious effects on social interaction parameters in WT controls. One week after cessation of treatment, beneficial effects of bumetanide were still detectable notably on the duration of nose contacts *(genotype x treatment: F_2,42_=9.3, p<0.0001)* and grooming episodes after social contact *(genotype x treatment: F_2,42_=16.4, p<0.0001)*, depending on the dose (Fig.1 C and S3).

In the 3-chamber test (Fig. 1D, more parameters in Fig. S2C), *Oprm1^-/-^* mice showed a severe impairment in social preference, as evidenced by equivalent number of nose contacts made with the mouse and the toy, and even longer nose contacts made with the toy over the mouse. Chronic NaBr completely restored social preference in mutant mice, which displayed more frequent (*genotype x treatment x stimulus: F_6,160_= 3.5, p<0.001*) and longer (*genotype x treatment x stimulus: F_6,160_= 8.9, p<0.0001*) nose contacts with the mouse since the lowest dose of bromide administered. This resulted in a normalization of their preference ratio from 10 mg/kg NaBr and over (*genotype x treatment: F_6,160_=11.9, p<0.0001*). In this test, chronic bumetanide at the lowest dose restored a preference for making more frequent nose contacts with the mouse over the toy (*genotype x treatment x stimulus: F_2,42_=17.7, p<0.0001).* Bumetanide treatment in mutant mice dose-dependently increased the duration of nose contacts with the mouse, but failed to reduce the duration of nose contacts with the toy (*genotype x treatment x stimulus: F_2,42_=32.5, p<0.0001)* leading to significant but partial recovery of social preference ratio in *Oprm1^-/-^* mice (*genotype x treatment: F_2,42_= 38.4, p<0.0001)*.

In conclusion, chronic but not acute NaBr treatment restored social behavior in *Oprm1^-/-^* mice in a dose-dependent manner, and these beneficial effects were superior to those of chronic bumetanide treatment.

### Sodium bromide reduced stereotypic behaviors and anxiety in *Oprm1^-/-^* mice

We next assessed the effects of chronic bromide on non-social behaviors in the same cohorts of *Oprm1^-/-^* mice (timeline in Fig. 1A, more parameters in Fig. S4). Regarding stereotypic behavior, *Oprm1^-/-^* mice displayed spontaneous stereotypic circling and head shakes (Fig. 2A) that were decreased under NaBr treatment since the lowest dose, more consistently for the former (*genotype x treatment: F_6,161_=4.6, p<0.001)* than for the latter (*genotype x treatment: F_6,161_=7.0, p<0.0001)*. In *Oprm1^+/+^* control mice, NaBr dose-dependently increased the number of grooming episodes (*genotype x treatment: F_6,161_=4.5, p<0.001)* and head shakes (*genotype x treatment: F_6,161_=7.0, p<0.0001)*; in both mouse lines, NaBr increased the number of rearing episodes in a dose-dependent manner (*treatment: F_6,161_=13.2, p<0.0001*). Under the same conditions, bumetanide suppressed circling and head shakes in mutant mice, and reduced the number of rearing episodes in both mouse lines. In the marble burying test (Fig. 2B), NaBr treatment did not suppressed excessive burying in mutant mice (*genotype: F_6,163_=13.6, p<0.001)* and globally increased burying (*treatment: F_6,163_=2.4, p<0.05).* Similarly, chronic bumetanide failed to suppress excessive marble burying in *Oprm1^-/-^* mice (*genotype: F_2,46_=14.7, p<0.001)*. In the Y-maze exploration test (Fig. 2C), bromide decreased the number of perseverative same arm returns in *Oprm1* null mice to wild-type levels since the dose of 30 mg/kg (*genotype x treatment: F_6,161_=6.7, p<0.0001)* while bumetanide globally failed to suppress those (*genotype: F_2,46_=19.7, p<0.001)*, despite a tendency for a decrease observed for the dose of 0.5 mg/kg.

**Fig. 2.**
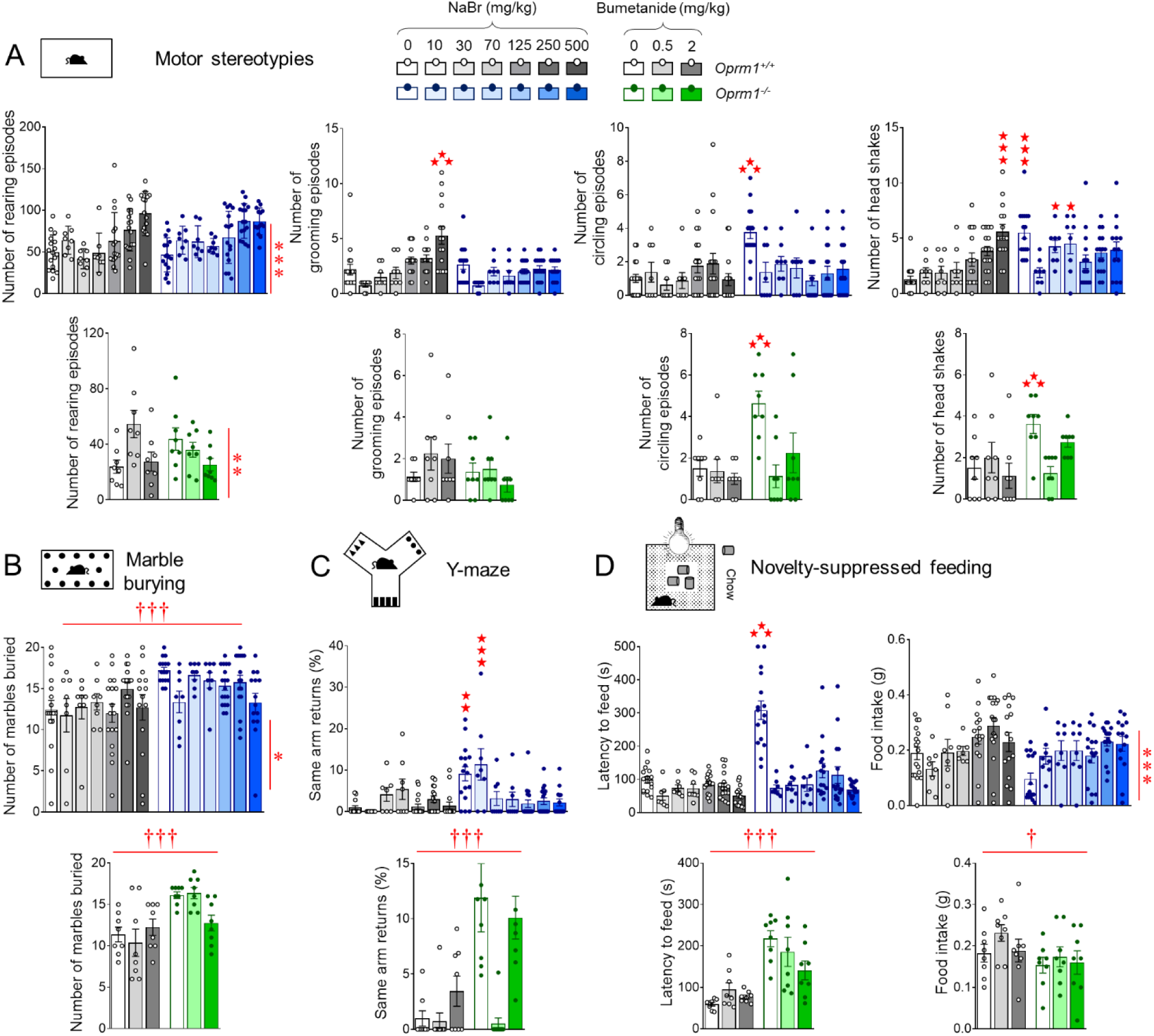
Chronic sodium bromide treatment reduced stereotypic behaviors and anxiety in *Oprm1^-/-^* mice. See timeline of experiments and animal numbers in Fig. 1A. (A) Chronic NaBr administration suppressed stereotypic circling episodes in *Oprm1^-/-^* mice since the dose of 10 mg/kg and less consistently reduced the number of head shakes (doses over 125 mg/kg). In wild-type controls, NaBr at 500 mg/kg increased the frequency of grooming episodes and head shakes. Bumetanide suppressed stereotypic circling and head shakes. (B) In the marble burying test, chronic bromide globally increased the number of buried marbles in both mouse lines; bumetanide failed to demonstrate significant effects. (C) In the Y-maze, NaBr suppressed perseverative same arm returns from the dose of 70 mg/kg; bumetanide had not significant effect despite an obvious tendency for the dose of 0.5 mg/kg to relieve perseveration. (D) In the novelty-suppressed feeding test, sodium bromide normalized the latency to feed in *Oprm1* null mice to wild-type levels since the lowest dose tested and increased food intake in all mice; bumetanide had no significant effect in this test. Results are shown as scatter plots and mean ± sem. Daggers: genotype effect, asterisks: treatment effect, solid stars: genotype x treatment interaction (comparison with wild-type vehicle condition) (two-way ANOVA followed by Newman-Keuls post-hoc test). One symbol: p<0.05, two symbols: p<0.01; three symbols: p<0.001. More behavioral parameters in Fig. S4.

We assessed anxiety levels in *Oprm1^-/-^* mice and their WT counterparts using the novelty suppressed feeding test (Fig. 2D). Mutant mice displayed exaggerated anxiety in this test, with increased latency to eat and reduced food intake once back in their home cage. Chronic bromide normalized eating latency (*genotype x treatment: F_6,160_=6.7, p<0.0001)* to wild-type levels in *Oprm1* knockout mice and increased food intake in both mouse lines (*treatment: F_6,160_=7.0, p<0.0001)* since the lowest dose tested. Chronic bumetanide had no detectable effect on either latency to eat (*genotype: F_2,42_=27.1, p<0.0001)* or food intake (*genotype: F_2,42_=5.6, p<0.05)* in this test.

Together, these results indicate that chronic bromide and bumetanide treatments both reduced stereotypic behavior in *Oprm1^-/-^* mice but only bromide treatment demonstrated anxiolytic effects.

In a next series of experiments, we verified that NaBr administered via oral route (oral gavage 4-5 days, 250 mg/kg once per day) relieved autistic-like deficits and motor stereotypies in *Oprm1^-/-^* mice in a similar way as it did after intra-peritoneal injection (Figure S5). Also, we assessed the behavioral effects of chronic administration of another bromide salt, KBr. A dose of KBr equivalent to 250 mg/kg NaBr being toxic in pilot experiments, we thus lowered the dose to 145 mg/kg KBr, equivalent to 125 mg/kg NaBr. Beneficial effects of NaBr treatment in *Oprm1^-/-^* mice were fully replicated, if not exceeded, by KBr in tests assessing social, repetitive and anxious behavior (Figure S6). Thus, therapeutic effects of NaBr or KBr in *Oprm1^-/-^* mice were attributable to bromide ions.

### Chronic sodium bromide relieved social behavior deficits, stereotypies and excessive anxiety in *Fmr1^-/-^* and *Shank3^Δex13-16-/-^* mice

We then questioned whether beneficial effects of NaBr on autistic-like symptoms may generalize to other mouse models of ASD, here the *Fmr1* null and *Shank3^Δex13-16^* knockout mouse lines. To this purpose, we evaluated the effects of chronic NaBr administration at the dose of 250 mg/kg on autism-sensitive behaviors in these lines (Fig. 3A, more parameters in Fig. S7 and S8).

**Fig. 3.**
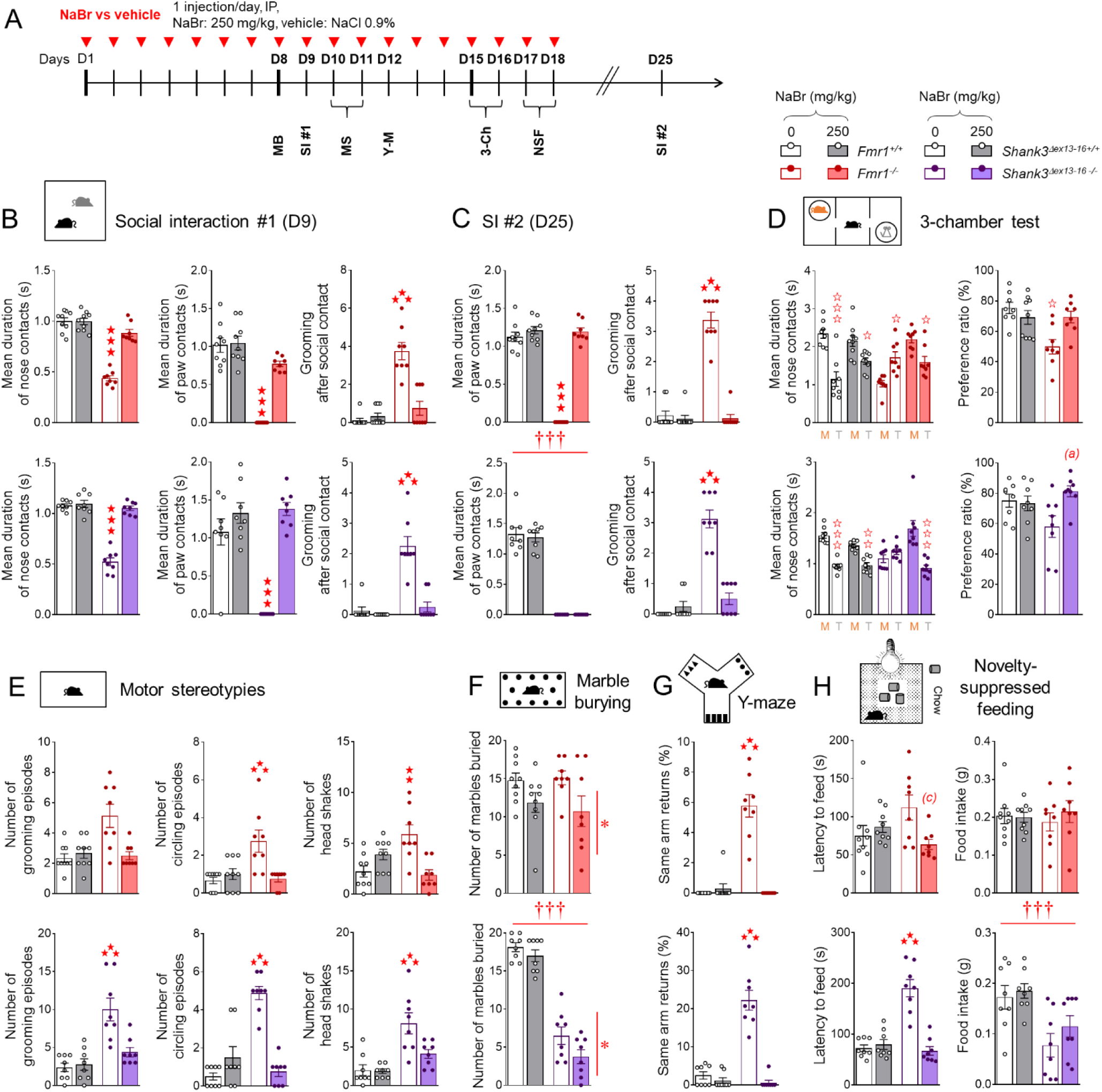
Chronic sodium bromide administration relieved social behavior deficits, stereotypies and exacerbated anxiety in *Fmr1^-/-^* and *Shank3^Δex13-16-/-^* mice. (A) *Fmr1^-/-^* or *Shank3^Δex13-16-/-^* and their respective wild-type counterparts were treated with NaBr (0 or 250 mg/kg; n=8 mice per genotype and treatment) once daily for 18 days. Behavioral testing started on D8; social interaction was retested 1 week (D25) after cessation of chronic administration. (B) In the direct social interaction test, chronic NaBr treatment normalized interaction parameters to wild-type levels in both *Fmr1* and *Shank3* mutant lines. (C) One week after cessation of treatment, these beneficial effects were fully maintained in *Fmr1^-/-^* mice while they were detected for some parameters only in *Shank3^Δex13-16-/-^* mice. (D) In the 3-chamber test, chronic NaBr administration rescued preference for making longer nose contacts with the mouse in *Shank3^Δex13-16-/-^* mice, resulting in increased preference ratio. (E) Chronic sodium bromide treatment suppressed stereotypic circling and head shakes in *Fmr1^-/-^* and *Shank3^Δex13-16-/-^* mice and normalized grooming in the latter. (F) NaBr reduced marble burying in *Fmr1^-/-^* and *Fmr1^+/+^* while it had no effect on reduced burying in *Shank3^Δex13-16-/-^* mice. (G) During Y-maze exploration, chronic NaBr suppressed perseverative same arm returns in both *Fmr1* and *Shank3* mutant lines. (H) Finally, in the novelty- suppressed feeding test, sodium bromide-treated *Fmr1^-/-^* or *Shank3^Δex13-16-/-^* mice displayed reduced or normalized latency to feed, respectively, but no modification in their food intake. Results are shown as scatter plots and mean ± sem. Daggers: genotype effect, solid stars: genotype x treatment interaction (comparison to wild-type vehicle condition), open stars: genotype x treatment x stimulus interaction (mouse versus object comparison), (a) genotype x treatment interaction (comparison with knockout vehicle condition, p<0.001), (c) genotype x treatment interaction (comparison to knockout vehicle condition, p<0.05) (two-way ANOVA or three-way ANOVA with stimulus as repeated measure, followed by Newman-Keuls post-hoc test). One symbol: p<0.05, two symbols: p<0.01; three symbols: p<0.001. More behavioral parameters in Fig. S7 and S8. 3-Ch: 3-chamber test, AAR: alternate arm returns, M: mouse, MB: marble burying, MS: motor stereotypies, NSF: novelty-suppressed feeding, SAR: same arm returns, SPA: spontaneous alternation, SI: social interaction, T: toy, Y-M: Y-maze.

As concerns social behavior, during a direct social interaction test (Fig. 3B), chronic bromide in *Fmr1^-/-^* as well as *Shank3^Δex13-16-/-^* mice restored the duration of nose *(genotype x treatment – Fmr1: F_1,30_=49.5, p<0.0001; Shank3^Δex13-16^: F_1,28_=73.0, p<0.0001)* and paw contacts *(genotype x treatment- Fmr1: F_1,30_=17.7, p<0.0001; Shank3^Δex13-16^: F_1,28_=23.2, p<0.0001)*, normalized the number of following episodes *(genotype x treatment - Fmr1: F_1,30_=13.7, p<0.0001; Shank3^Δex13-16^: F_1,28_=11.8, p<0.0001)* and suppressed grooming after social contact *(genotype x treatment- Fmr1: F_1,30_=30.1, p<0.0001; Shank3^Δex13-16^: F_1,28_=25.0, p<0.0001)*. One week after interruption of NaBr treatment (Fig. 3C), significant beneficial effects were still detected on the duration of paw contacts *(genotype x treatment: F_1,30_=132.7, p<0.0001)* and number of grooming episodes after a social contact *(genotype x treatment: F_1,30_=87.0, p<0.0001)* in *Fmr1* null mice. In *Shank3^Δex13-16^* mice, no effect of previous bromide treatment was longer detected on the duration of paw contacts (*genotype: F_1,28_=388.2, p<0.0001)*, whereas grooming after social contact remained efficiently suppressed (*genotype x treatment: F_1,28_=55.3, p<0.0001)*.

We further assessed social behavior under chronic bromide exposure using the 3- chamber test (Fig. 3D). Although *Fmr1* knockout mice made more frequent nose contacts with the mouse versus the object in this test, they spent as much time in contact with the living mouse as with the object and made longer nose contacts with the object, which demonstrates disrupted social preference. Chronic NaBr treatment restored a preference for spending more time exploring the mouse (*genotype x treatment x stimulus: F_1,29_= 4.5, p<0.05*) and making longer nose contacts with the mouse versus the object (*genotype x treatment x stimulus: F_1,29_= 18.4, p<0.001)* in *Fmr1* mutants, which normalized their preference ratio (*genotype x treatment: F_1,29_=5.2, p<0.05)* without modifying the number of nose contacts they made with either stimulus (*stimulus*: *F_1,29_=21.4, p<0.0001)*. Likewise, *Shank3^Δex13-16-/-^* mice treated with vehicle failed to spend more time with the mouse over the toy in this test; they made more frequent nose contacts with the mouse but of equivalent duration with both stimuli. In contrast, mutants treated with NaBr spent more time in contact with the mouse (*genotype x treatment x stimulus: F_1,28_= 4.5, p<0.05*) and made longer nose contacts with their congener versus the toy (*genotype x treatment x stimulus: F_1,28_= 26.9, p<0.0001)*, resulting in increased preference ratio (*genotype x treatment: F_1,28_=5.9, p<0.05)* with no change in the number of nose contacts (*stimulus: F_1,28_=37.4, p<0.0001).* Thus, chronic NaBr administration rescued social behavior deficits in *Fmr1* null and *Shank3^Δex13-16^* knockout mice.

As regards stereotypic behavior, *Fmr1^-/-^* and *Shank3^Δex13-16-/-^* mice displayed more frequent spontaneous grooming (significant in the latter only), circling episodes and head shakes than WT controls (Fig. 3E). Chronic NaBr treatment normalized all these parameters to WT levels (*Fmr1 – circling, genotype x treatment: F_1,30_=11.9, p<0.01; head shakes, genotype x treatment: F_1,30_=68.0, p<0.001; Shank3^Δex13-16^ – grooming, genotype x treatment: F_1,28_=10.2, p<0.01, circling, genotype x treatment: F_1,28_=48.4, p<0.0001; head shakes, genotype x treatment: F_1,28_=5.3, p<0.05).* In the marble burying test (Fig. 3F), chronic NaBr reduced the number of buried marbles in both *Fmr1^+/+^* and *Fmr1^-/-^* mice (*treatment effect: F_1,30_=7.3, p<0.05)*; *Shank3^Δex13-16-/-^* mice displayed a severe deficit in marble burying (*genotype effect: F_1,28_=198.7, p<0.*0001) and bromide treatment decreased burying in both mutant and WT mice (*treatment effect: F_1,28_=4.8, p<0.05).* In the Y-maze (Fig. 3G), *Fmr1* null and *Shank3^Δex13-16^* knockout mice displayed more frequent perseverative same arm returns that were suppressed under chronic bromide (*Fmr1* - *genotype x treatment: F_1,30_=61.9, p<0.0001; Shank3^Δex13-16^ - genotype x treatment: F_1,28_=47.5, p<0.0001).* Thus, chronic NaBr treatment reduced stereotypic and perseverative behaviors in *Fmr1^-/-^* and*Shank3^Δex13-16-/-^* mice.

As regards anxiety, a tendency for *Fmr1* null mice for increased latency to eat in the novelty-suppressed feeding test (Fig. 3H) did not reach significance; however chronic bromide reduced this latency (*genotype x treatment: F_1,30_=6.7, p<0.05)*. *Shank3^Δex13- 16-/-^* mice took significantly longer to eat in the center of the arena and NaBr administration normalized this latency to WT levels (*genotype x treatment: F_1,28_=35.0, p<0.0001)*. Bromide treatment had no effect on food intake, reduced in *Shank3^Δex13-16^* knockout mice (*genotype: F_1,28_=15.6, p<0.001).* Therefore, chronic NaBr administration demonstrated anxiolytic properties in *Fmr1^-/-^* and *Shank3^Δex13-16-/-^* mice.

### Chronic sodium bromide modulates transcription in the reward circuit of *Oprm1^-**/-**^* **mice**

To shed light on the molecular mechanism involved in beneficial effects of chronic sodium bromide administration, we assessed the effects of a 2-week NaBr treatment on gene expression in *Oprm1* null mice across five regions of the brain reward/social circuit: NAc, CPu, VP/Tu, MeA and VTA/SNc. Mice underwent a first session of social interaction after one week under treatment and a second 45 min before sacrifice for qRT-PCR experiment (Fig. 4A). We focused primarily on genes coding for chloride transporters (*Slc12a2* [NKCC1], *Slc12a4,5,6,7* [KCC1,2,3,4, respectively], *ClCa1*), GABAA receptor subunits (*Gabra1,2,3,4,5, Gabrb1,2*) and glutamate receptors (*Grm2,4,5)* and subunits *(Grin2a,2b*). In addition, we evaluated the expression of marker genes of neuronal expression and plasticity (*Fos*, *Bdnf*), social behavior (*Oxt*) and striatal projection neurons (SPNs; *Crh*, *Drd1a*, *Drd2*, *Htr6, Pdyn*, *Penk*).

**Fig. 4.**
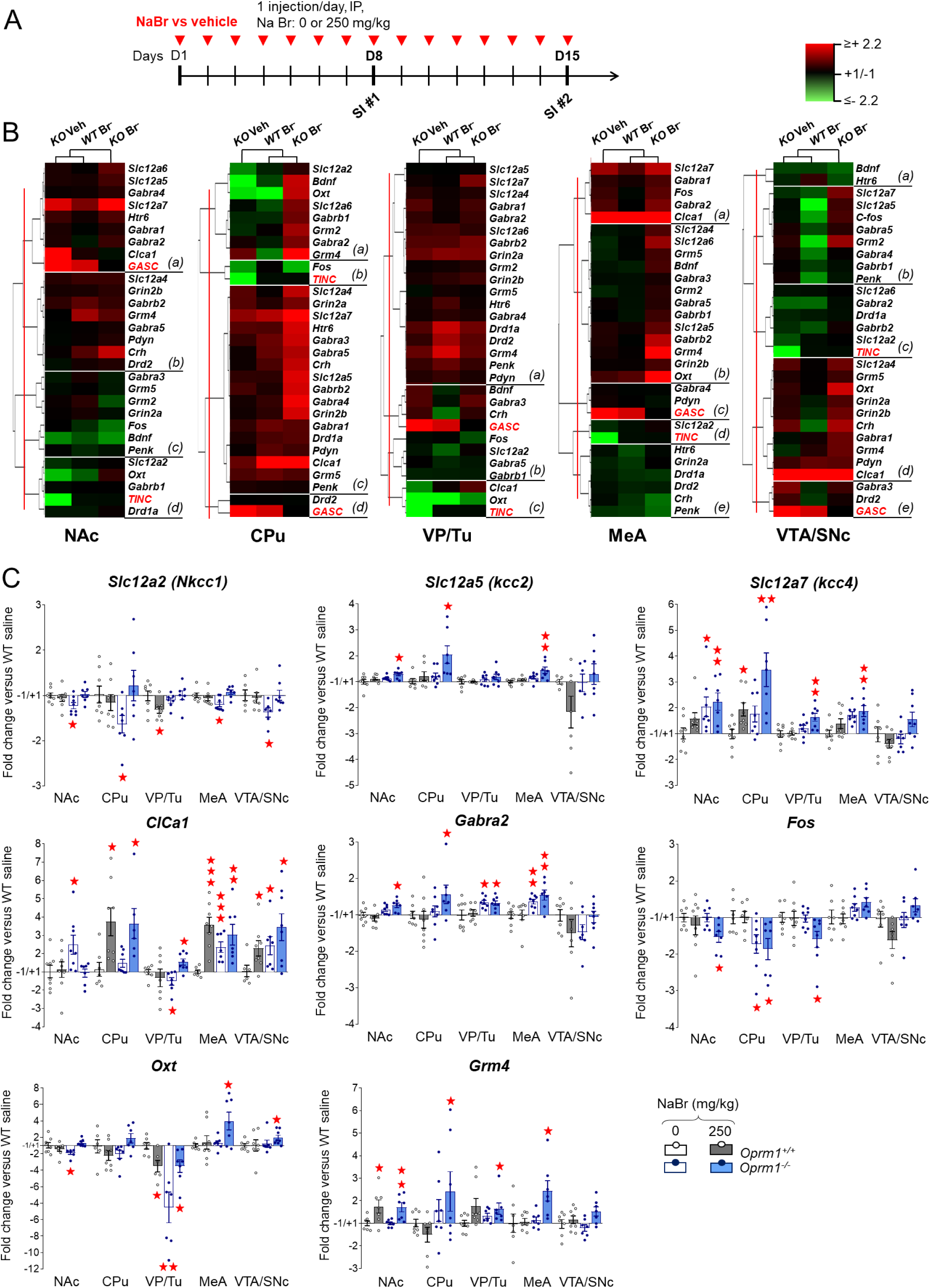
Chronic sodium bromide treatment induced transcriptional modifications in the reward circuit of *Oprm1^-/-^* mice. (A) *Oprm1^+/+^* and *Oprm1^-/-^* mice were treated for 2 weeks with either vehicle or NaBr (250 mg/kg, i.p. once a day). They undergone two sessions of direct social interaction, at D8 and D15, the latter 45 min before sacrifice for qRT-PCR experiment. We evaluated the expression of 27 genes of interest in 5 regions of the reward/social circuit: the NAc, CPu, VP/Tu, MeA and VTA/SNc. (B) Clustering analysis revealed that the most contrasted transcriptional profiles were observed in *Oprm1^-/-^* mice under NaBr versus vehicle treatment; mRNA levels, however, were poorly correlated with behavioral parameters (red characters, GASC: grooming after social contact, TINC: time in nose contact). Bromide treatment induced rather than normalized gene expression in *Oprm1* null mice, most remarkably in the CPu. (C) Focusing on candidate genes, we unraveled a global upregulation of chloride transporters, notably *Slc12a2*, *Slc12a5* and *Slc12a7*, coding respectively for NKCC1, KCC2 and KCC4, and *Clca1*, coding for the calcium-activated chloride channel regulator 1 (CLCA1) and GABA-related genes, among which *Gabra2*, coding for the α2 subunit of the GABAA receptor, in mutant mice under NaBr treatment. In contrast, *Fos* expression was reduced in the basal ganglia of NaBr-treated *Oprm1^-/-^* mice while *Bdnf* expression was up-regulated in the CPu and MeA. Finally, chronic NaBr in mutant mice increased the levels of *Oxt* and *Grm4*, coding respectively for oxytocin and mGlu4 receptor, across brain regions. Gene (n=8 per group) expression data are expressed as fold change versus *Oprm1^+/+^* - vehicle group (scatter plots and mean ± SEM). Comparison to *Oprm1^+/+^* - vehicle group (two-tailed t-test): One star p<0.05, two stars p<0.01, three stars p<0.001. qRT-PCR data used for clustering are displayed in Table S2.

We performed hierarchical clustering analysis of qRT-PCR data for each brain region to visualize the influence of NaBr treatment on gene expression (Fig. 4B, Table S2). Overall, transcriptional profiles in *Oprm1^-/-^* mice under vehicle and NaBr treatment differed the most, while mRNA levels correlated poorly with social interaction parameters (Fig. S9). These results indicate that bromide induced transcriptional changes on its own rather than it normalized gene expression in *Oprm1* knockouts (as observed for behavioral parameters). This was particularly true in the CPu, where *Oprm1^-/-^* mice under bromide treatment displayed predominant up-regulated gene expression (clusters a and c).

This overall profile was confirmed when focusing on candidate genes (Fig. 4C). We ought to acknowledge here that the sample number of mice allocated to each experimental condition was low to address the complex influences of genotype and pharmacological treatment, which may have limited the statistical power. For this reason, we focused our attention on gene expression regulations affecting either several brain regions for the same gene, or several genes of the same family. Strikingly, chronic NaBr administration increased the expression of all tested chloride transporters in *Oprm1^-/-^* mice. Indeed, the expression of *Slc12a2* was decreased in all brain regions but the VP/Tu of mutant mice, and normalized under bromide treatment. Chronic NaBr up-regulated the expression of *Slc12a5* and *Slc12a7* in the NAc, CPu and MeA of *Oprm1* null mice for the former, together with the VP/Tu for the latter. In mutant mice, *ClCa1* mRNA levels were increased in the NAc, MeA and VTA/SNc while reduced in the VP/Tu; they were normalized by NaBr treatment in the NAc, increased in the VP/Tu and maintained high in the MeA and VTA/SNc. In the CPu, bromide increased *ClCa1* transcription in mice of both genotypes. As regards the GABAergic system, chronic NaBr in *Oprm1^-/-^* mice stimulated the expression of *Gabra2*, coding for the α2 subunit of the GABAA receptor, in the NAc and CPu and left this expression high in the VP/TU and MeA. Remarkably, bromide consistently upregulated the expression of *Gabra3, Gabra4, Gabra5, Gabrg1* and *Gabrb2* in the CPu of mutant mice (Table S2). Bromide treatment down-regulated the expression of the early gene *Fos* in the NAc and VP/Tu and maintained it low in the CPu of *Oprm1* knockouts. *Oxt* (coding for oxytocin) mRNA levels were decreased in the NAc and VP/Tu of *Oprm1^-/-^* mice and normalized by NaBr in the former and partially in the latter; bromide induced *Oxt* expression in the MeA and VTA/SNc. Finally, chronic NaBr upregulated the expression of *Grm4*, coding for the metabolic glutamate receptor mGlu4, in all brain regions but the VTA/SNc of mutant mice, and in the NAc of wild-type controls. Transcriptional results thus indicate that bromide administration had a major impact on the expression of Cl^-^ transporters; meanwhile it also regulated the expression of several key players of the GABA system, marker genes of neuronal activity and plasticity, and genes more specifically involved in the control of social behavior.

### Synergistic effects of chronic bromide and mGlu4 receptor facilitation in *Oprm1* null mice

Intrigued by the increase in *Grm4* transcription in *Oprm1^-/-^* mice under bromide treatment, whose autistic-like symptoms were relieved when stimulating mGlu4 activity (*51*), we then addressed the question of a potential shared mechanism of action between these treatments. To this aim, we studied the effects of combined administration of chronic liminal doses of NaBr (70 mg/kg) and VU0155041 (1 mg/kg) in *Oprm1^-/-^* and *Oprm1^+/+^* mice (Fig. 5A, more parameters in Fig. S10).

**Fig. 5.**
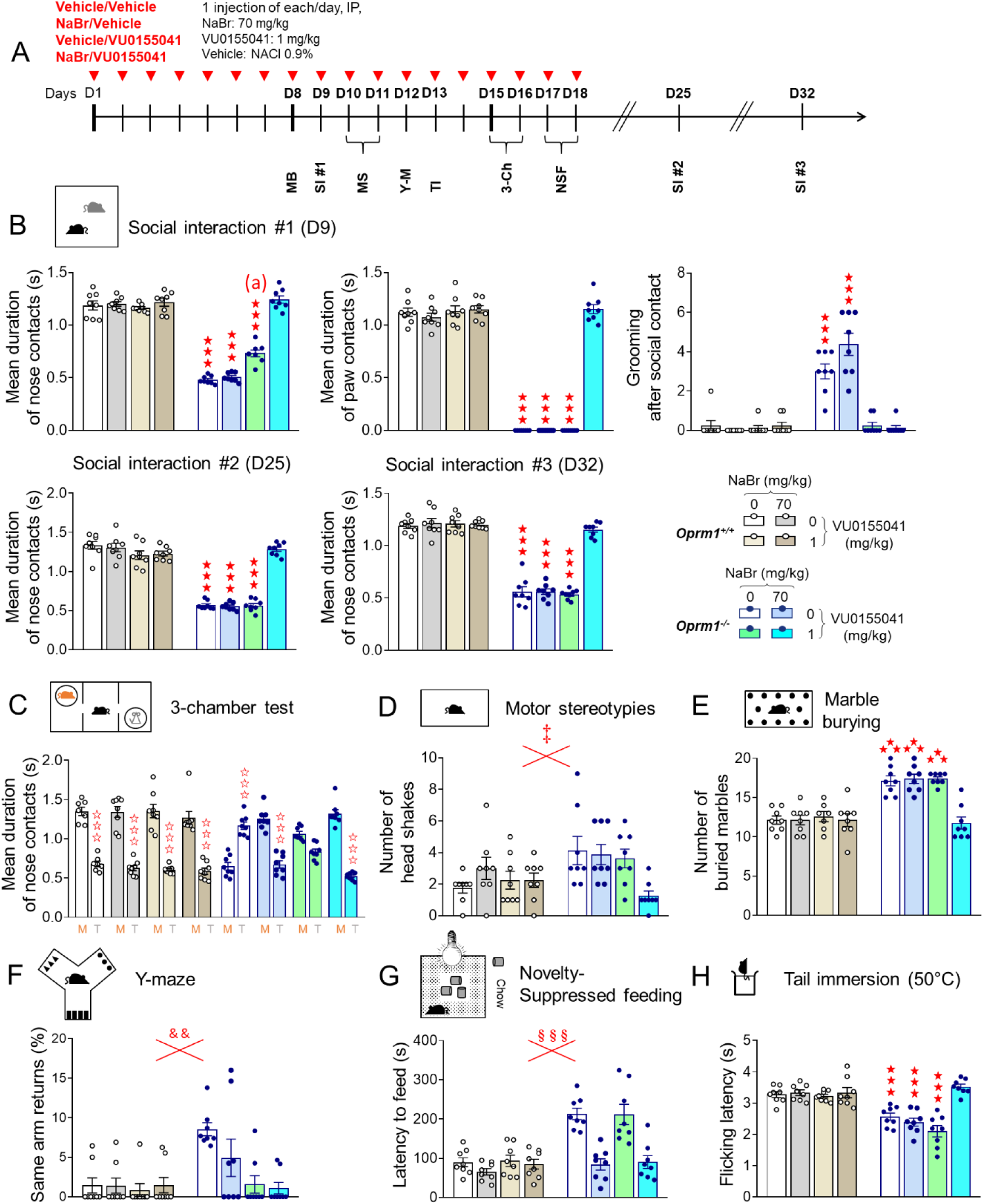
Beneficial effects of sodium bromide and VU0155041, a positive allosteric modulator of mGlu4 receptors, were synergistic in *Oprm1^-/-^* mice. (A) *Oprm1^+/+^* and *Oprm1^-/-^* mice were treated either with vehicle, NaBr (70 mg/kg), VU0155041 (1 mg/kg) or NaBr and VU0155041 (70 and 1 mg/k, respectively; 8 mice per genotype and dose) once daily for 18 days. Behavioral testing started on D8; social interaction was retested 1 week and 2 weeks after cessation of chronic administration. (B) In the direct social interaction test, NaBr and VU0155041 treatments demonstrated synergistic effects in restoring the duration of nose and paw contacts in *Oprm1^-/-^* mice. VU0155041 at 1 mg/kg, however, was sufficient to suppress grooming after social contact. Beneficial effects of combined NaBr/VU0155041 were fully maintained one and two weeks after cessation of treatment. (C) In the 3-chamber test, VU0155041 at 1 mg/kg increased the duration of nose contacts with the mouse to that of the toy in *Oprm1^-/-^* mice; NaBr at 70 mg/kg and combined NaBr/VU0155041 treatment fully restored longer nose contacts with the mouse. (D) Combined NaBr/VU0155041 administration reduced head shakes in *Oprm1^-/-^* and *Oprm1^+/+^* mice and (E) normalized marble burying in *Oprm1* null mice only. (F) VU0155041 treatment was sufficient to suppress perseverative same arm returns in *Oprm1^-/-^* mice exploring the Y-maze and (G) NaBr administration was sufficient to normalize their latency to feed in novelty-suppressed feeding test. (H) In the tail immersion test at 50°C, only combined NaBr/VU0155041 treatment restored flicking latency of *Oprm1^-/-^* mice to wild-type levels. Results are shown as scatter plots and mean ± sem. Solid stars: genotype x NaBr x VU0155041 interaction (comparison to wild-type vehicle condition), open stars: genotype x stimulus x NaBr x VU0155041 interaction (mouse versus object comparison), (a) genotype x NaBr x VU0155041 interaction (comparison with knockout vehicle condition, p<0.001), double dagger: NaBr x VU0155041 interaction, ampersand: genotype x VU0155041 interaction, section: genotype x NaBr interaction (three-way or four-way ANOVA followed by Newman-Keuls post-hoc test). One symbol: p<0.05, two symbols: p<0.01; three symbols: p<0.001. 3-Ch: 3-chamber test, M: mouse, MB: marble burying, MS: motor stereotypies, NSF: novelty-suppressed feeding, SI: social interaction, T: toy, TI: tail immersion, Y-M: Y-maze. More behavioral parameters in Fig. S10 and S11.

In the direct social interaction test (Fig. 5B), NaBr at 70 mg/kg had no detectable effects on behavioral parameters in *Oprm1^-/-^* mice (see Fig. 1B); VU0155041 partially rescued the duration of their nose contacts and suppressed grooming episodes after a social contact. When the two treatments were combined, social interactions parameters in mutant mice were normalized to wild-type levels, as illustrated by rescued duration of nose contacts (*genotype x bromide x VU0155041: F_1,56_=31.1, p<0.0001*) or paw contacts (*genotype x bromide x VU0155041: F_1,56_=140.3, p<0.0001*) and normalized number of grooming episodes after social contact (*genotype x bromide x VU0155041: F_1,56_=5.8, p<0.05*). Restoration of the mean duration of nose contacts was fully preserved one (*genotype x bromide x VU0155041: F_1,56_=40.9, p<0.0001*) and two (*genotype x bromide x VU0155041: F_1,56_=57.3, p<0.0001*) weeks after cessation of treatment. In the 3-chamber test, NaBr at 70 mg/kg was sufficient to restore longer nose contacts with the mouse versus the toy in *Oprm1^-/-^* mice (*genotype x NaBr x stimulus: F_1,55_=61.6, p<0.0001*) (Fig. 5C).

As regards stereotypic behavior, combined NaBr and VU0155041 treatments reduced the number of head shakes in *Oprm1^-/-^* and *Oprm1^+/+^* mice (*bromide x VU0155041: F_1,56_=4.1, p<0.05*) (Fig. 5D) and normalized marble burying in mutants (*genotype x NaBr x VU0155041: F_1,56_=9.5, p<0.001*) (Fig. 5E). Chronic VU0155041 was sufficient to suppress perseverative same arm returns during Y-maze exploration in *Oprm1* null mice (*genotype x VU0155041: F_1,56_=11.3, p<0.01)* (Fig. 5F). In the novelty-suppressed feeding test, chronic NaBr was similarly sufficient to normalize latency to feed (*genotype x VU0155041: F_1,56_=11.3, p<0.01)* (Fig. 5G). Finally, having in mind that nociceptive thresholds are lowered in *Oprm1* null mice, we tested the effects of bromide and VU0155041 on this parameter. At 50°C, combined NaBr and VU0155041 treatment normalized flicking latency, while each compound given alone was ineffective (*genotype x NaBr x VU0155041: F_1,56_=20.5, p<0.0001*) (Fig. 5H). Together, these results indicate that bromide administration and facilitation of mGlu4 activity exert synergistic beneficial effects on autistic-like behavior in *Oprm1^-/-^* mice.

### Bromide ions behave as positive allosteric modulators of the mGlu4 glutamate receptor

Chloride ions have been shown to facilitate mGlu4 signaling (*56*). Here we assessed whether synergistic *in vivo* effects of bromide treatment and VU0155041 administration may result from a modulation of mGlu4 activity by bromide ions, in addition to their ability to trigger an upregulation of *Grm4* expression (see Fig. 4).

We measured mGlu4 signaling under glutamate stimulation in HEK293T cells transiently expressing mGlu4 receptors and Gαqi9, a Gi/Gq chimeric G-protein (*57*) that allows mGlu4 to activate the phosphoinositide pathway. Receptor activation was then evaluated by measuring intracellular Ca^2+^ release or inositol monophosphate (IP1, Fig. 6A). The experiments were performed in buffers containing either a physiological concentration of chloride ions (100 mM, supplemented with 50 mM gluconate to maintain equivalent osmolarity between mediums), 150 mM of chloride ions (classical buffer for cell culture studies) or 100 mM of chloride ions and 50 mM of bromide ions, to compare the effects of modulating chloride and bromide concentrations within a physiological range on mGlu4 activity.

**Fig. 6.**
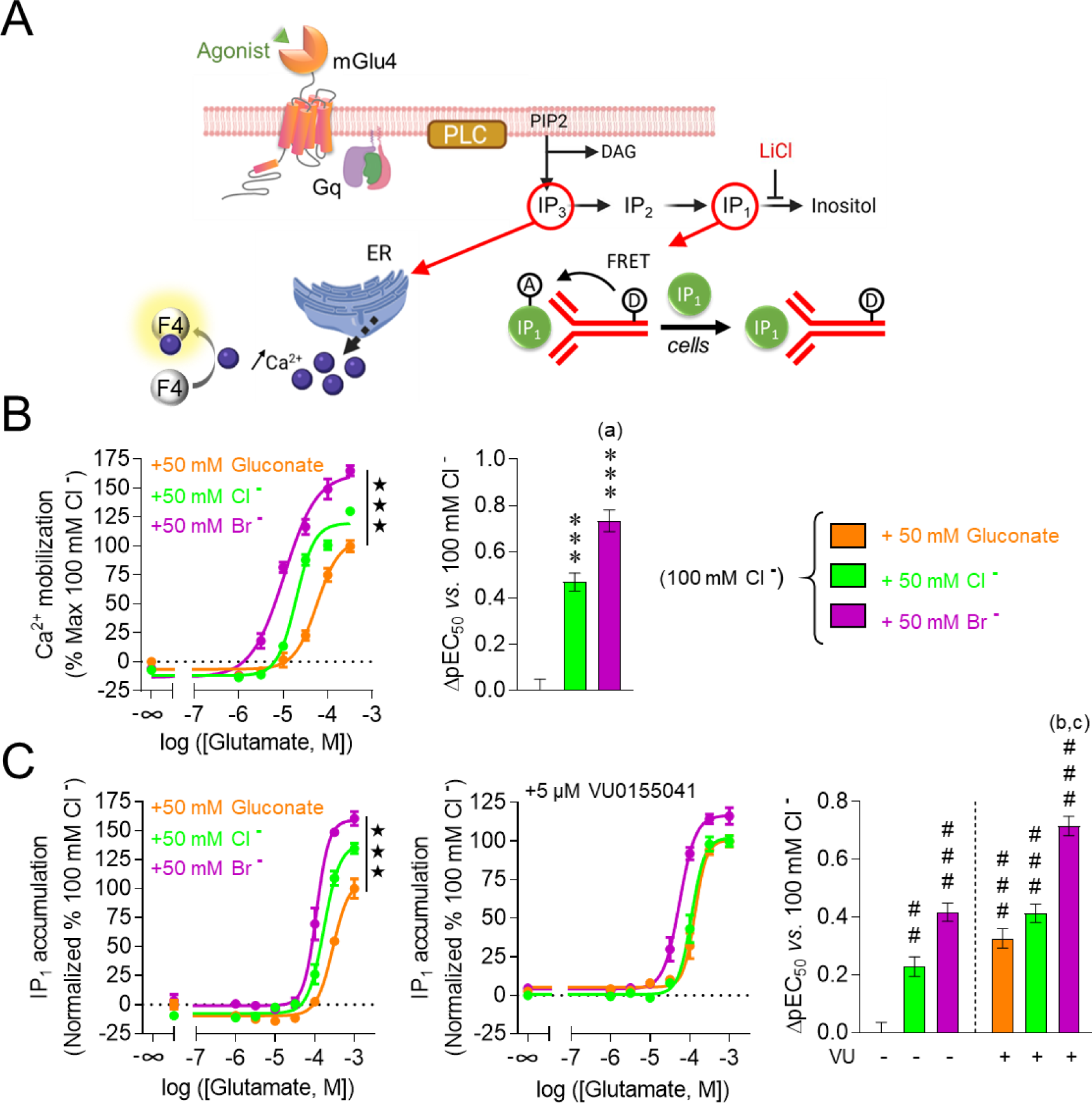
Bromide ions behave as PAMs of the mGlu4 receptor and shows synergistic effects with the mGlu4 PAM VU0155041. (A) Signaling cascade of mGlu4 receptor when coupled with the chimeric G-protein Gαqi9 (to allow the recruitment of the phosphoinositide pathway) and experimental principles of the calcium mobilization (Panel B) and IP1 accumulation assays (Panel C). (B) In the calcium mobilization assay, bromide ions behave as PAM of mGlu4, demonstrating broader effects than chloride ions on both pEC50 and Emax. (C) In the IP1 accumulation assay, bromide confirmed its PAM effects; when supplemented with VU0155041, facilitation of mGlu4 signaling was even increased, as seen by a further rise in ΔpEC50. Results are shown as mean ± SEM of three independent experiments realized in triplicates. Solid stars: effect of ion concentration on Emax; asterisks: effect of ion concentration on ΔpEC50, comparison with physiological conditions (100 mM Cl^-^ + 50 mM gluconate); (a): comparison with high chloride condition (150 mM Cl^-^, p<0.0001); hashtags: effect of ion concentration and VU0155041 on ΔpEC50, comparison with physiological conditions (100 mM Cl^-^ + 50 mM gluconate); (b) and (c): comparison with bromide conditions (100 mM Cl^-^ + 50 mM Br-) with or without VU0155041 (p<0.0001) (One-way ANOVA, followed by Tukey’s post-hoc test). Two symbols: p<0.001, three symbols: p<0.0001. DAG: diacylglycerol; ER: endoplasmic reticulum; F4: Fluo4 calcium probe; FRET: fluorescence resonance energy transfer; PLC: phospholipase C; IP1/2/3: inositol mono/di/triphosphate.

Compared with physiological concentration of chloride (100 mM Cl^-^ + 50 mM gluconate), addition of 50 nM bromide significantly improved glutamate potency, showing a 0.73 ± 0.05 log increase in pEC50 (left panel; *ion concentration: F_2,6_=104.2, p<0.0001*) in the Ca^2+^ assay (Fig. 6B). Compared with an equivalent concentration of chloride (+ 50 mM Cl^-^), bromide showed a higher efficacy (ΔpEC50: 0.27±0.04) (right panel; *ion concentration: F_2,6_=66.7, p<0.0001*). Further, bromide increased glutamate efficacy, with a 65 ± 9% rise in maximal mGlu4-triggered calcium release (Emax, left panel; *ion concentration: F_2,21_=56.2, p<0.0001*).

When measuring IP1 production (Fig. 6C), bromide increased glutamate efficacy (Emax, maximal IP1 production) within a similar range (60 ± 10%) as observed when measuring Ca^2+^ release (left panel; *ion concentration: F_2,24_=22.1, p<0.0001*). Despite technical limitations (less amplification at this step of the signaling cascade), pEC50 was consistently increased in presence of bromide ions (left panel; *ion concentration: F_2,6_=104.2, p<0.0001*; ΔpEC50 between physiological Cl^-^ concentration and after addition of 50 nM Br^-^: 0.42 ± 0.03) though to a lower extend as those measured in the Ca^2+^ assay. When 5µM of VU0155041 were added, bromide significantly increased glutamate potency compared with physiological concentration of chloride (ΔpEC50: 0.72 ± 0.03). Bromide and VU0155041 combination was also more effective in increasing glutamate potency than VU0155041 (ΔpEC50: 0.39 ± 0.03) or bromide (ΔpEC50: 0.30 ± 0.03) alone (*ion concentration and VU0155041*: *F_5,12_=49.4, p<0.0001)*. In conclusion, bromide ions behaved as PAMs of the mGlu4 receptor in heterologous cells. These PAM effects were superior to those of chloride ions, and synergized with those of VU0155041. Together, these results provide a molecular mechanism for synergistic effects of bromide and VU0155041 in *Oprm1* null mice, and suggest that benefits of bromide treatment in mouse models of ASD involved, at least in part, a facilitation of mGlu4 activity.

## DISCUSSION

In the current study, we tested whether bromide ions would relieve autistic-like symptoms in mouse models of ASD. We first assessed bromide effects in the *Oprm1^-/-^* mouse model of ASD, which recapitulates a wide range of autistic-like core and secondary symptoms, including severe deficits in social interaction and communication (*51, 58–61*), stereotypic behavior, exacerbated anxiety (*51, 60*) and increased susceptibility to seizures (*51, 62*) but displays preserved cognitive performance (*52*). In this model, chronic bromide dose-dependently rescued social behavior. NaBr treatment improved direct social interaction from the dose of 125 mg/kg, with complete rescue at 250 mg/kg. These doses are in the lower range of acute anticonvulsant of bromide in rodents (0.2 to 2 g/kg) (*63, 64*) and slightly superior to chronic KBr antiepileptic doses in dogs (20-100 mg/kg/day; initial dose up to 500 mg/kg) (*65, 66*) and humans (30-100 mg/kg/day) (*67, 68*). In the 3-chamber test, beneficial effects of chronic NaBr were detected since the dose of 10 mg/kg. In this test, however, social interactions occur only when the experimental mouse investigates its congener, with limited reciprocity. In contrast, social contacts during dyadic interaction are reciprocal, which can be stressful for each protagonist, justifying the long use of this test to measure anxiety levels (*69*). Yet, these data suggest that beneficial effects of NaBr on social behavior can be reached since the dose of 10 mg/kg.

Compared with chronic treatment, acute administration of NaBr had only minor effects on social interaction in *Oprm1* null mice, showing that bromide needed to accumulate to reach therapeutic efficacy, as observed in the clinics (*68*). Consistent with this, blood concentrations of bromide ions are known to progressively increase upon chronic treatment (*70, 71*). As its clearance is slow, bromide has a rather long half-life, about 8-14 days in human adults (*68, 72*). This pharmacokinetic profile may account for the carry-over effects of NaBr treatment detected in *Oprm1^-/-^* mice. NaBr also reduced stereotypic and perseverative behaviors in *Oprm1* knockouts, since the dose of 10-30 mg/kg, indicating benefit on the second diagnostic criteria for ASD. To date, no approved treatment ameliorates both diagnostic dimensions of autism. Behavioral improvements were replicated after repeated oral administration of NaBr, in agreement with previous reports of oral bioavailability (*72*). Further, bromide treatment, known as anxiolytic (*45*), consistently decreased anxiety levels in *Oprm1^-/-^* mice, a significant advantage as anxiety disorders are among the most frequent comorbidities in ASD (*3, 5*). Finally, beneficial effects of chronic NaBr administration in *Oprm1^-/-^* mice were replicated using KBr, long used as an antiepileptic drug in human (*47*) and veterinary (*65*) medicine, despite its narrow therapeutic window due to cardiotoxicity of potassium ions (K^+^) (*73*). Convergent effects of NaBr and KBr salts demonstrate that bromide ions are indeed the active compound to relieve ASD-like symptoms.

Effects of chronic NaBr administration were compared to those of the NKCC1 inhibitor bumetanide. Although the most promising drug developed based on the E/I balance hypothesis, bumetanide has been scarcely tested in animal models of ASD. We observed partial beneficial effects of chronic bumetanide on autistic-like behavior in *Oprm1^-/-^* mice. In contrast, previous studies have reported a complete rescue of autistic-like social deficits following bumetanide treatment in animal models of ASD, but under conditions where bumetanide was administered preventively rather than curatively. Indeed, bumetanide restored social communication in two rodent models of ASD, rats exposed to valproate *in utero* and *Fmr1* knockout mice, when their mothers were treated with bumetanide during pregnancy (*40*). Similarly, repeated bumetanide administration rescued social preference when used to prevent early-life seizures in rats (*74*). In the present study, bumetanide was given in adult *Oprm1^-/-^* mice, when neurodevelopmental deficits were fully installed, which may account for partial improvements. These effects were consistent with clinical reports of beneficial effects of the NKCC1 antagonist on social abilities in children and adults with ASD diagnosis (*41, 42*), maybe too subtle however to be confirmed in larger cohorts (*43*), whilst this treatment efficiently reduced stereotypic behavior (*43*). Importantly, such consistency confers predictive value to the *Oprm1* null mouse model, boding well for clinical translation of bromide effects.

After comprehensive testing in *Oprm1* null mice, we extended our observation of beneficial effects of bromide administration to two additional mouse models, *Fmr1^-/-^* and *Shank3^Δex13-16-/-^* mice, with longer-lasting effects in the former. Interestingly, the three models of ASD used in this study differed in their genetic cause and E/I profile. As regards excitatory neurotransmission, *Fmr1* mutants display more frequent miniature excitatory postsynaptic currents (mEPSCs) in the hippocampus (*40*), persistent activity states in the somatosensory cortex (*75*) and excessive mGluR1/5- dependent signaling (*75, 76*), suggesting a global facilitation of glutamatergic transmission. Mutations in *Shank3* lead to reduced frequency and amplitude of mEPSCs in the striatum (*55, 77*) and hippocampus (*78*) and reduced long-term potentiation (LTP) in these two structures (*79, 80*); in *Oprm1^-/-^* mice, hippocampal LTP is decreased (*52*) and the morphology of asymmetrical synapses is altered in the striatum while the expression of key glutamate-related genes is reduced (*51*). These data point to impaired glutamatergic neurotransmission in the latter models. On the inhibition side, multiple alterations of the GABAergic system have been documented in *Fmr1* null mice (*81, 82*), with reduced expression of GABAA receptor subunits (*27, 30, 82*) and consistent impairment in GABAergic transmission (*30, 83*). Such deficit, however, is not observed in the striatum, where GABA release is increased (*84*), possibly a consequence of impaired endocannabinoid-mediated long-term depression (LTD) that represses the activity of D2 dopamine receptor-expressing striatal projection neurons (D2-SPNs) (*53*). Remarkably, endocannabinoid-mediated LTD is also deficient in *Shank3^Δex13-16^* mice, and consequent excessive D2-SPN tone has been linked to stereotyped behavior (*85*). In *Oprm1^-/-^* mice, facilitating the activity of mGlu4 receptor, which is known to repress D2-SPN activity (*86*), relieved autistic-like behavior (*51*). Together, these data designate GABAergic D2-SPNs as a potential common target for bromide effects on the E/I balance across mouse models.

Transcriptional effects of chronic bromide treatment in *Oprm1^-/-^* versus *Oprm1^+/+^* mice are consistent with an influence on the E/I balance. In the absence of NaBr treatment, *Oprm1* null mice displayed low levels of *Slc12a2* transcripts (NKCC1), suggesting that Cl^-^ gradient was altered. Strikingly, NaBr administration restored or increased the expression of all genes coding for Cl^-^ co-transporters tested in these mice, the importer NKCC1 as well as the exporters KCC2-4 and CLCA1 (a modulator of the calcium-activated Cl^-^ channel Anoctamine-1/TMEM16A (*87*)). All these transporters have been related to autistic-like features in mouse models (*40, 88, 89*), putting a spell on chloride homeostasis as a core neurobiological mechanism in ASD. Increased expression of Cl^-^ transporters under chronic NaBr treatment may have been a regulatory response triggered by Br^-^ accumulation, to maintain the extracellular/intracellular gradient of anion concentrations. Cl^-^ and Br^-^ sharing similar physicochemical properties, this increase likely favored a high turnover of both anion species and, thus, depolarizing effects of GABA (*90*). Moreover, chronic bromide increased the transcription of genes coding for several GABAA receptor subunits. Previous studies in the field of epilepsy have evidenced a facilitating influence of bromide on GABAergic inhibition (*48, 91*). Together, these data suggest that NaBr treatment increased GABA-mediated inhibition. Accordingly, Fos expression was decreased in the basal ganglia of *Orpm1* null mice following social interaction, a likely consequence of facilitated GABAergic inhibition.

Besides effects on E/I balance, bromide treatment in *Oprm1^-/-^* mice stimulated the expression of *Oxt*, coding for the social neuropeptide oxytocin, notably in the NAc, where it plays a key role in modulating social reward (*92*). We observed similar restoration in *Oprm1^-/-^* mice after that they undergone an appetitive social conditioning (*60*). Moreover, oxytocin administration rescued social ultrasonic vocalizations in *Oprm1* null mice (*93*). Thus, restored *Oxt* mRNA levels in mutants may have contributed to behavioral improvements under bromide administration. Finally, although *Grm4* mRNA levels were not low in the striatum of *Oprm1^-/-^* mice as in previous studies (*51, 60*), maybe due to different experimental conditions, chronic NaBr increased its expression in most brain regions studied. By facilitating mGlu4 expression, bromide may thus have relieved autistic-like symptoms, as observed under facilitation of mGlu4 activity by VU0155041 treatment (*51, 94*). We challenged this hypothesis by co-administering liminal doses of bromide and VU0155041 to *Oprm1^-/-^* mice, and revealed a major synergistic effect. Moreover, we evidenced that, in heterologous cells, bromide ions behave as better mGlu4 PAMs than it was previously shown for chloride ions (*56*). This finding suggests that beneficial effects of Br^-^ in ASD models involve a facilitation of mGlu4 signaling. Moreover, the PAM effect of bromide synergizes with that of VU0155041, providing a molecular mechanism for synergy *in vivo*. Yet, this mechanism may not have been the only one. At cellular level, VU0155041 represses the activity of D2-SPNs (*86*), found disinhibited in *Fmr1^-/-^* and *Shank3^Δex13-16-/-^* mice, while bromide promotes GABAergic transmission. Thus, synergistic action of bromide and mGlu4 PAM may also lie in their common inhibition of a neuronal population, D2-SPNs, a hypothesis that will deserve further investigation. Hence, bromide ions act at multiple levels, on chloride gradient, GABA-mediated neurotransmission and mGlu4 signaling, which all converge towards neuronal inhibition, and this may account for its large spectrum of beneficial effects in mouse models of ASD (Figure 7).

**Fig. 7.**
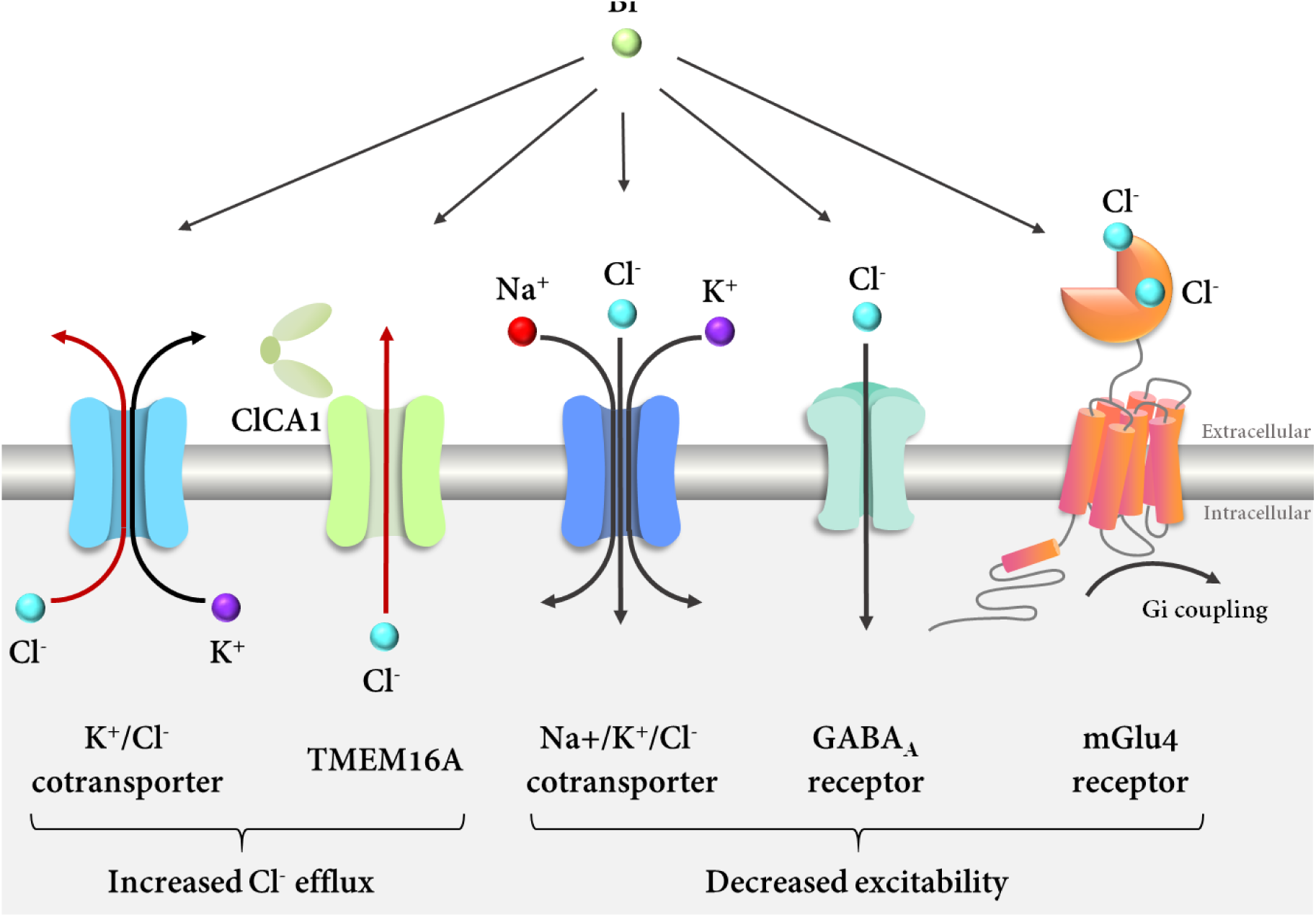
Different cellular mechanisms by which bromide ions may modulate E/I balance and neurotransmission in the autistic brain. At transcriptional level, chronic bromide treatment up- regulated the expression of genes coding for Cl^-^ extruders KCCs and the secreted activator of TMEM16A, ClCA1, in *Oprm1^-/-^* mice. By such mechanism, bromide ions may have increased Cl^-^ efflux (dark red), making Cl^-^ more available within the extracellular compartment. In parallel, bromide was found to induce the expression of genes coding for NKCC1, GABAA and the mGlu4 receptor in *Oprm1* mutants. Interestingly, increased extracellular Cl^-^ concentrations facilitate the activation of these three molecular actors, which converges towards decreased neuronal excitability. All previous mechanisms would thus concur to increase neuronal inhibition. Intriguingly, due to its physicochemical properties, Br^-^ can substitute to Cl^-^ at all levels; by binding to mGlu4, it behaves as a positive allosteric modulator of this receptor, even more potent than Cl^-^. ClCa1 and TMEM16A representations were adapted from (*87*) and Cl^-^ location on mGlu4 from (*56*).

We ought to consider here some limitations of this study. The low sample number of mice from both sexes allocated to each genotype and treatment group resulted in low statistical power to examine the complex interactions between genotype and bromide effects in our transcriptional study. Further investigations will be required to explore the molecular mechanisms underlying beneficial effects of bromide in mouse models of ASD. Also, we tested the effects of bromide treatment in adult mice whilst autism is diagnosed early in life and early behavioral/cognitive interventions have proven to be more efficient in relieving autistic features (*95, 96*). In future studies, we will administer bromide at young age in mouse models of ASD to delineate the optimal developmental window for its therapeutic action. Whether it would not be verified in youngsters, though, therapeutic benefit of bromide treatment would remain a major breakthrough in the field of autism, which affects a large and highly vulnerable population of adults worldwide (*97*).

In conclusion, the present study reports the therapeutic potential of chronic bromide treatment, alone or in combination with a PAM of mGlu4 receptor, to relieve core symptoms of ASD. Beneficial effects of bromide were observed in three mouse models of ASD with different genetic causes, supporting high translational value. Moreover, bromide has a long history of medical use, meaning that its pharmacodynamics and toxicity are well known, which, combined with long lasting effects as well as excellent oral bioavailability and brain penetrance, are strong advantages for repurposing.

## MATERIALS AND METHODS

### Key resources table

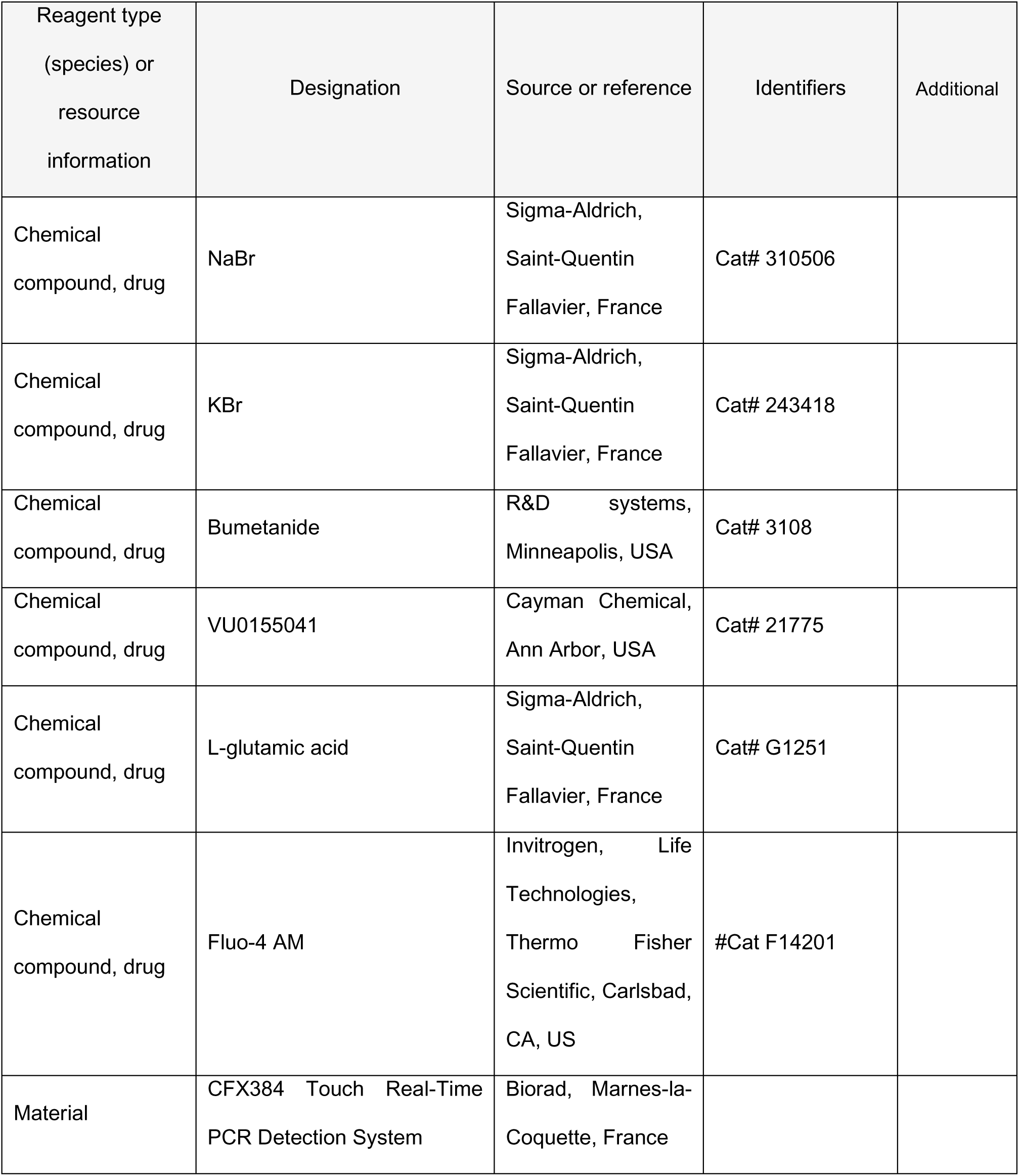
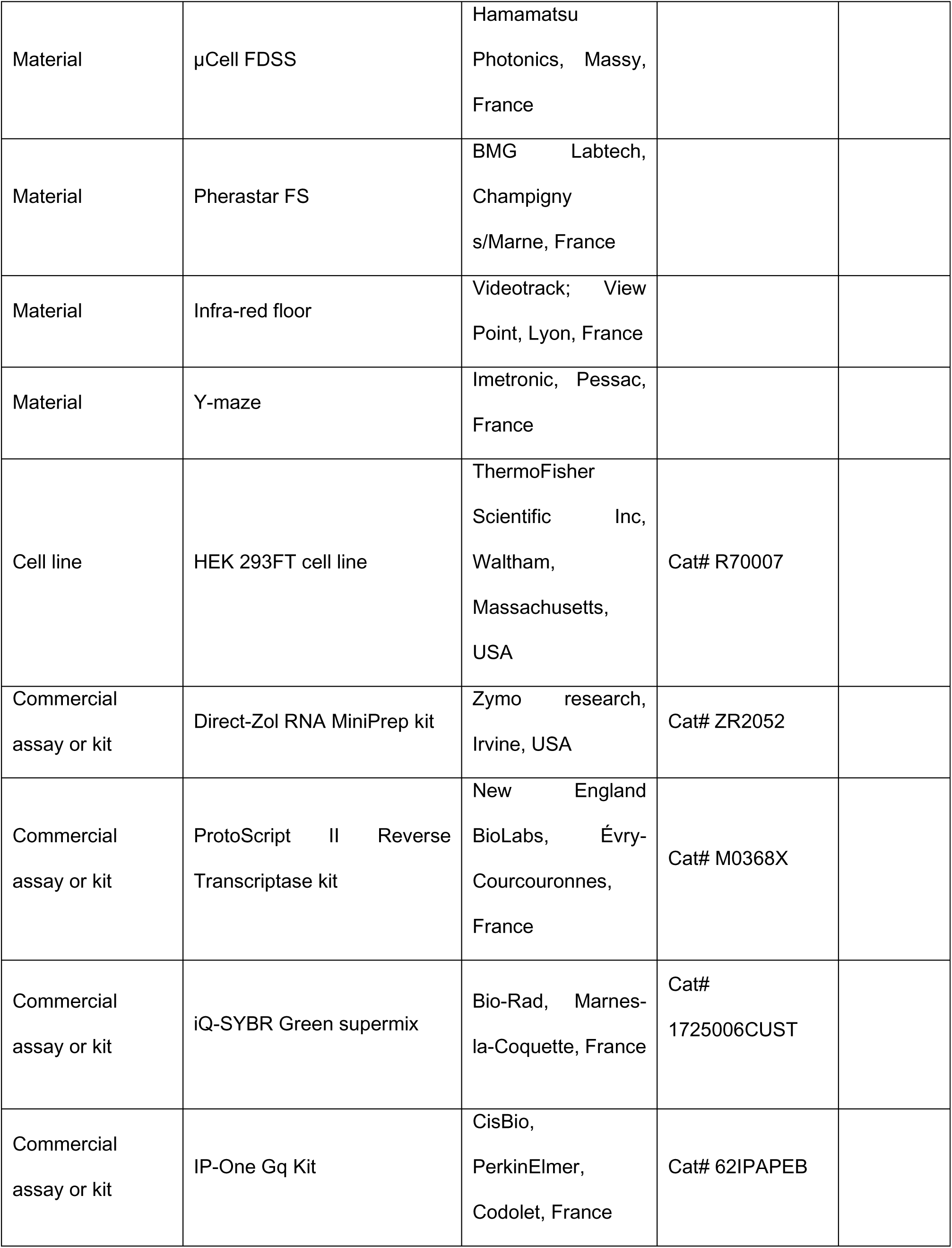
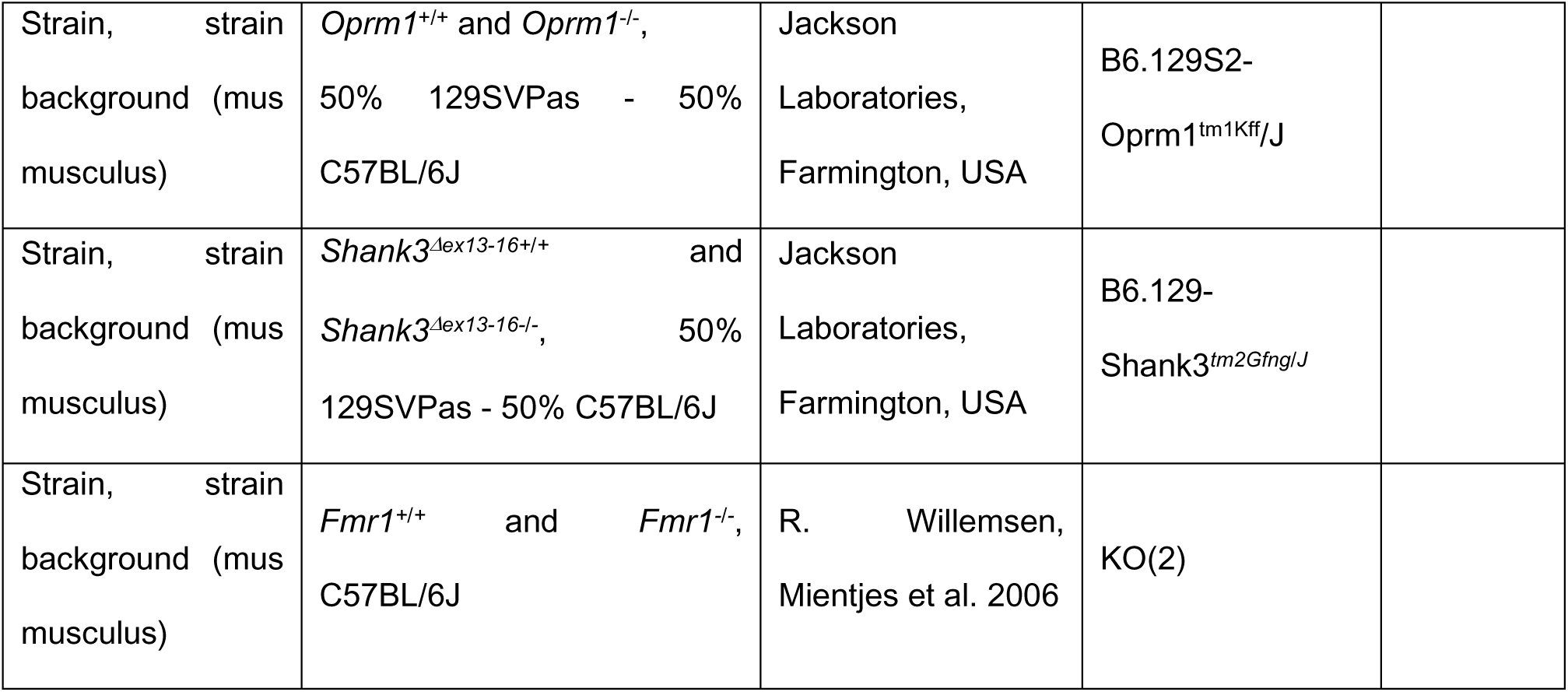

### Animals, breeding procedures and housing conditions

The *Oprm1^-/-^* (B6.129S2-Oprm1^tm1Kff^/J) (*98*) and *Shank3^Δex13-16-/-^* (B6.129- Shank3*^tm2Gfng/J^*, so called *Shank3B^-/-^*, lacking the PDZ domain) (*55*) mouse lines were acquired from Jackson Laboratories (Farmington, USA) and bred on a hybrid background: 50% 129SVPas - 50% C57BL/6J. *Fmr1*-KO(2) mice (*99*) were generously provided by R. Willemsen (Erasmus University Medical Center, Rotterdam, The Netherlands) and bred on a C57BL/6J background. Equivalent numbers of male and female mice were generated in-house from homozygous parents, bred from heterozygous animals, to prevent genetic derivation. This breeding scheme favored social deficits in mutant mice by maintaining them together during early post-natal development. Except otherwise stated, animals were group-housed and maintained on a 12hr light/dark cycle (lights on at 7:00 AM) at controlled temperature (21±1°C); food and water were available *ad libitum*. Experiments were analyzed blind to genotypes and experimental condition. All experimental procedures were conducted in accordance with the European Communities Council Directive 2010/63/EU and approved by the Comité d’Ethique en Expérimentation animale Val de Loire (C2EA- 19).

### Drugs

Mice were treated with vehicle (NaCl 0.9%; ip, 10 or 20 ml/kg), NaBr (Sigma-Aldrich, Saint-Quentin Fallavier, France) administered either chronically (once a day, 10, 30, 70, 125, 250 and 500 mg/kg; i.p. or per os, in a volume of 20 ml/kg – except for combination with VU0155041: 10 mg/kg) or acutely (250 mg/kg, 20 ml/kg), KBr (Sigma-Aldrich, Saint-Quentin Fallavier, France; 145 mg/kg; i.p., 20 ml/kg), bumetanide (R&D systems, Minneapolis, USA, 0.5 and 2 mg/kg; i.p., 20 ml/kg) or VU0155041 (Cayman Chemical, Ann Arbor, USA, once a day, i.p., 1 mg/kg, 10 ml/kg). Doses of bumetanide were chosen based on previous studies in rodent models of ASD (*40, 74*); liminal dose of VU0155041 was set based on our previous studies (*51, 100*) and a pilot experiment showing no detectable effect in the social interaction test. When treatment was given chronically, behavioral testing started 8 days after beginning of daily administration. Treatment was maintained for 8 to 18 consecutive days (see timelines in Fig. 1, 4 and 5), allowing thorough behavioral phenotyping. On testing days, or when treatment was given acutely, drugs (or vehicle) were administered 30 min before behavioral assays.

### Behavioral experiments

When assessing effects of chronic administration, experiments were performed successively (timelines in Fig. 1, 4 and 5) (*51, 100*). Testing order was chosen to minimize the incidence of anxiety on later assays. Direct social interaction and novelty suppressed feeding were performed in 4 equal square arenas (open fields, 50 x 50 cm) separated by 35cm-high opaque grey Plexiglas walls over a white Plexiglas platform (View Point, Lyon, France). Stimulus mice used for the three-chamber test were 8-14-week-old grouped-housed male or female wild-type mice, socially naive to the experimental animals.

#### Social abilities

##### Direct social interaction test

On testing day, a pair of unfamiliar mice (not cage mates, age-, sex- and treatment-matched) was introduced in each arena for 10 min (15 lx). Each arena received a black plastic floor (transparent to infrared). The total amount of time spent in nose contact (nose-to-nose, nose-to-body or nose-to-anogenital region), the number of these contacts, the time spent in paw contact and the number of these contacts, grooming episodes (allogrooming), notably ones occurring immediately (<5s) after a social contact, as well as the number of following episodes were scored *a posteriori* on video recordings (infrared light-sensitive video camera) using an ethological keyboard (Labwatcher®, View Point, Lyon, France) by trained experimenters and individually for each animal (*51, 100*). The mean duration of nose and paw contacts was calculated from previous data (*101–103*).

##### Three-chamber social preference test

The test apparatus consisted of a transparent acrylic box (exterior walls blinded with black plastic film); partitions divided the box into three equal chambers (40 x 20 x 22.5 cm). Two sliding doors (8 x 5 cm) allowed transitions between chambers. Cylindrical wire cages (18 x 9 cm, 0.5 cm diameter-rods spaced 1 cm apart) were used to contain the mouse interactor and object (soft- toy mouse). The test was performed in low-light conditions (15 lx) to minor anxiety. Stimulus wild-type mice were habituated to confinement in wire cages for 2 days before the test (20 min/day). On testing day, the experimental animal was introduced to the middle chamber and allowed to explore the whole apparatus for a 10-min habituation phase (wire cages empty) after the sliding doors were raised. The experimental mouse was then confined back in the middle-chamber while the experimenter introduced an unfamiliar wild type age and sex-matched animal into a wire cage in one of the side-chambers and a soft toy mouse (8 x 10 cm) in the second wire cage as a control for novelty. Then the experimental mouse was allowed to explore the apparatus for a 10- min interaction phase. The time spent in each chamber, the time spent in nose contact with each wire cage (empty: habituation; containing a mouse or a toy: interaction), as well as the number of these nose contacts were scored *a posteriori* on video recordings using an ethological keyboard (Labwatcher®, View Point, Lyon, France) by trained experimenters. The mean duration of nose contacts was calculated from these data (*101–103*). The relative position of stimulus mice (versus toy) was counterbalanced between groups.

#### Stereotyped behaviors

##### Motor stereotypies

To detect motor stereotypies in mutant versus wild-type animals, mice were individually placed in clear standard home cages (21×11×17 cm) filled with 3-cm deep fresh sawdust for 10 min (*104*). Light intensity was set at 30 lux. Trained experimenters scored numbers of head shakes, as well as rearing, burying, grooming, circling episodes and total time spent burying by direct observation.

##### Y-maze exploration

Spontaneous alternation behavior was used to assess perseverative behavior (*105–107*). Each Y-maze (Imetronic, Pessac, France) consisted of three connected Plexiglas arms (15×15×17 cm) covered with distinct wall patterns (15 lx). Floors were covered with lightly sprayed fresh sawdust to limit anxiety. Each mouse was placed at the center of a maze and allowed to freely explore this environment for 5 min. The pattern of entries into each arm was quoted on video-recordings. Spontaneous alternations (SPA), i.e. successive entries into each arm forming overlapping triplet sets, alternate arm returns (AAR) and same arm returns (SAR) were scored, and the percentage of SPA, AAR and SAR was calculated as following: total / (total arm entries -2) * 100.

##### Marble-burying

Marble burying was used as a measure of perseverative behavior (*108*). Mice were introduced individually in transparent cages (21×11×17 cm) containing 20 glass marbles (diameter: 1.5 cm) evenly spaced on 4-cm deep fresh sawdust. To prevent escapes, each cage was covered with a filtering lid. Light intensity in the room was set at 40 lux. The animals were removed from the cages after 15 min, and the number of marbles buried more than half in sawdust was quoted.

#### Anxiety-like behavior

##### Novelty-suppressed feeding

Novelty-suppressed feeding (NSF) was measured in 24-hr food-deprived mice, isolated in a standard housing cage for 30 min before individual testing. Three pellets of ordinary lab chow were placed on a white tissue in the center of each arena, lit at 60 lx. Each mouse was placed in a corner of an arena and allowed to explore for a maximum of 15 min. Latency to feed was measured as the time necessary to bite a food pellet. Immediately after an eating event, the mouse was transferred back to home cage (free from cage-mates) and allowed to feed on lab chow for 5 min. Food consumption in the home cage was measured.

##### Nociceptive thresholds

Tail-immersion test. Nociceptive thresholds were assessed by immersing the tail of the mice (5 cm from the tip) successively into water baths at 48°C, 50°C and 52°C. The latency to withdraw the tail was measured at each temperature, with a cutoff of 10 s.

### Real-time quantitative PCR analysis

Brains were removed and placed into a brain matrix (ASI Instruments, Warren, MI, USA). Nucleus accumbens (NAc), caudate putamen (CPu), ventral pallidum/olfactory tubercle (VP/Tu), medial nucleus of the amygdala (MeA) and ventral tegmental area/substancia nigra pars compacta (VTA/SNc) were punched out/dissected from 1mm-thick slices (see Fig. S1). Tissues were immediately frozen on dry ice and kept at -80°C until use. For each structure of interest, genotype and condition, samples were processed individually (n=8). RNA was extracted and purified using the Direct-Zol RNA MiniPrep kit (Zymo research, Irvine, USA). cDNA was synthetized using the ProtoScript II Reverse Transcriptase kit (New England BioLabs, Évry-Courcouronnes, France). qRT-PCR was performed in quadruplets on a CFX384 Touch Real-Time PCR Detection System (Biorad, Marnes-la-Coquette, France) using iQ-SYBR Green supermix (Bio-Rad) kit with 0.25 µl cDNA in a 12 µl final volume in Hard-Shell Thin-Wall 384-Well Skirted PCR Plates (Bio-rad). Gene-specific primers were designed using Primer3 software to obtain a 100- to 150-bp product; sequences are displayed in Table S1. Relative expression ratios were normalized to the level of actin and the 2^−ΔΔCt^ method was applied to evaluate differential expression level. Gene expression values differing of the mean by more than two standard deviations were considered as outliers and excluded from further calculations.

### Cell culture and transfection

Human embryonic kidney (HEK) 293 cells were transiently transfected with rat mGlu4 receptor by electroporation together with a chimeric Gi/Gq protein to allow phospholipase activation and EAAC1, a glutamate transporter, to avoid influence of extracellular glutamate. For calcium mobilization, cells were seeded in a PLO-coated, black-walled, clear-bottomed, 96-well plate (Greiner Bio-One) at the density of 100,000 cells per well and for IPOne assay in a PLO-coated black 96-well plate (Greiner Bio-One) at the density of 50,000 cells per well. Cells were cultured in DMEM (Gibco^TM^, Life Technologies), supplemented with 10% fetal bovine serum. Medium was changed by GlutaMAX (Gibco^TM^, Life Technologies) to reduce extracellular glutamate concentration 3 h before experiment.

### Chloride and bromide buffers for *in vitro* assays

Chloride and bromide ion concentrations in the buffers used for *in vitro* experiments were chosen for best matching physiological conditions. It was previously shown that the total amount of halogen (chloride and bromide ions) in the cerebrospinal fluid and serum can reach 120-130 mM (*109, 110*) and that bromide is able to substitute up to 30% of chloride concentration (*111*), leading to a theoretical concentration of 91 mM chloride and 36 mM bromide. We thus chose the dose of 100 mM chloride as physiological control (complemented with gluconate NaC6H11O7 to maintain equivalent osmolarity between buffers (*56*)), and added or not 50 mM chloride or bromide to assess their effects *in vitro*. Chemicals were purchased from Sigma-Aldrich (Merck, L’lsle D’Abeau Chesnes, France). The pH of all buffers was adjusted to 7.4 before experiments.

For calcium mobilization assay, buffers with 100 mM NaCl, 2.6 mM KCl, 1.18 mM MgSO4, 10 mM D-glucose, 10 mM 4-(2-hydroxyethyl)-1-piperazineethanesulfonic acid (HEPES), 1mM CaCl2 and 0.5% (w/v) bovine serum albumin were used and supplemented with 50 mM of either NaCl, NaBr or NaC6H11O7 to reach 154.6 mM total anion concentration.

For IP1 accumulation assay, buffers with 46 mM NaCl, 4.2 mM KCl, 0.5 mM MgCl2, 10 mM HEPES, 1mM CaCl2, 50 mM NaC6H11O7 and 50 mM LiCl (to avoid IP1 degradation) were used and supplemented with 50 mM of either NaCl, NaBr or NaC6H11O7 to reach 203.2 mM total anion concentration.

### Calcium mobilization and IP1 accumulation assays

Calcium mobilization assay: 24 h after transfection, cells were loaded with 1 µM calcium-sensitive fluorescent dye (Fluo-4 AM; Invitrogen, Life Technologies) diluted in fresh Cl^-^ buffer (154.6 mM) for 1 h at 37°C and 5% CO2. Then, cells were washed and maintained in appropriate buffer supplemented with 4 mM probenecid. Agonists were also diluted in appropriate buffers. Ca^2+^ release was determined using µCell FDSS (Hamamatsu Photonics). Fluoresence was recorded for 60s at 480 nm excitation and 540 nm emission after 20 s record of baseline.

IP1 accumulation assay: Inositol monophosphate accumulation was determined using the IP-One HTRF (Homogenous Time Resolved Fluorescence) kit (Cisbio Bioassays, Perkin Elmer, Codolet, France) according to the manufacturer’s recommendations (*112*). Briefly, cells were stimulated to induce IP1 accumulation while being treated with test compounds in appropriate buffer for 30 min, at 37 °C and 5% CO2 before d2- labeled IP1 and Tb-labeled anti-IP1 antibody addition. After 1h incubation at RT, Pherastar FS (BMG Labtech) was used to record 620 nm and 665 nm emission after 337 nm excitation.

### Data analysis and statistics

#### In vivo experiments

Statistical analyses were performed using Statistica 9.0 software (StatSoft, Maisons-Alfort, France). For all comparisons, values of p<0.05 were considered as significant. Statistical significance in behavioral experiments was assessed using one or two-way analysis of variance (drug, stimulus and treatment effects) followed by Newman-Keuls post-hoc test. Significance of quantitative real-time PCR (qRT-PCR) results was assessed after transformation using a two-tailed t-test, as previously described (*100*). Unsupervised clustering analysis was performed on transformed qRT-PCR data using complete linkage with correlation distance (Pearson correlation) for genotype and treatment (Cluster 3.0 and Treeview software) (*51, 100*). When used for clustering analysis (Figure 7B), behavioral data were normalized to vehicle-vehicle condition and transformed using the same formula as qRT-PCR data.

#### In vitro experiments

Data were analyzed using Prism 6 software (GraphPad Software, San Diego, CA, USA). For IP1 accumulation assay, each HTRF-ratio was transformed in IP1 concentration using calibration curves for each buffer and then normalized against Mock-transfected cells to avoid buffer composition effect on HTRF signal. In all experiments, a 4-parameter concentration-response curve equation was used to fit data, and potency (EC50) was estimated as logarithms (log EC50). For clarity purpose, absolute (positive) logarithms (pEC50) were used. Emax represents the maximum response obtained at saturating agonist concentration. Data shown in the figures represent the means ± SEM of at least 3 experiments realized in triplicates. Statistical differences between pEC50, ΔpEC50 and Emax were determined using a one-way analysis of variance followed by Tukey’s post-test.

### Data availability

All data that support the findings of this study are available from the corresponding author upon request.

## Acknowledgments

We thank Dr Thierry Plouvier for inspiring initial discussions on this project, Pr Frédérique Bonnet-Brilhault for critical reading of the manuscript, Yannick Corde for technical support and Drs. Jorge Gandía and Sébastien Roux for assistance in performing behavioral experiments. We thank the Experimental Unit PAO-1297 (EU0028, Animal Physiology Experimental Facility, DOI: 10.15454/1.5573896321728955E12) from the INRAE-Val de Loire Centre for animal breeding and care.

## Author contributions

Conceptualization: CD, JK, JPP, JLM, JAJB; Methodology: JK, JPP, JLM, JAJB; Investigation: CD, AL, AB, DJ, JLM, JAJB; Visualization: CD, AB, JLM, JAJB; Funding acquisition: JPP, JLM, JAJB; Project administration: JK, JPP, JLM, JAJB; Supervision: JK, JPP, JLM, JAJB; Writing: CD, JLM, JAJB

## Supplementary Figures

**Figure S1.**
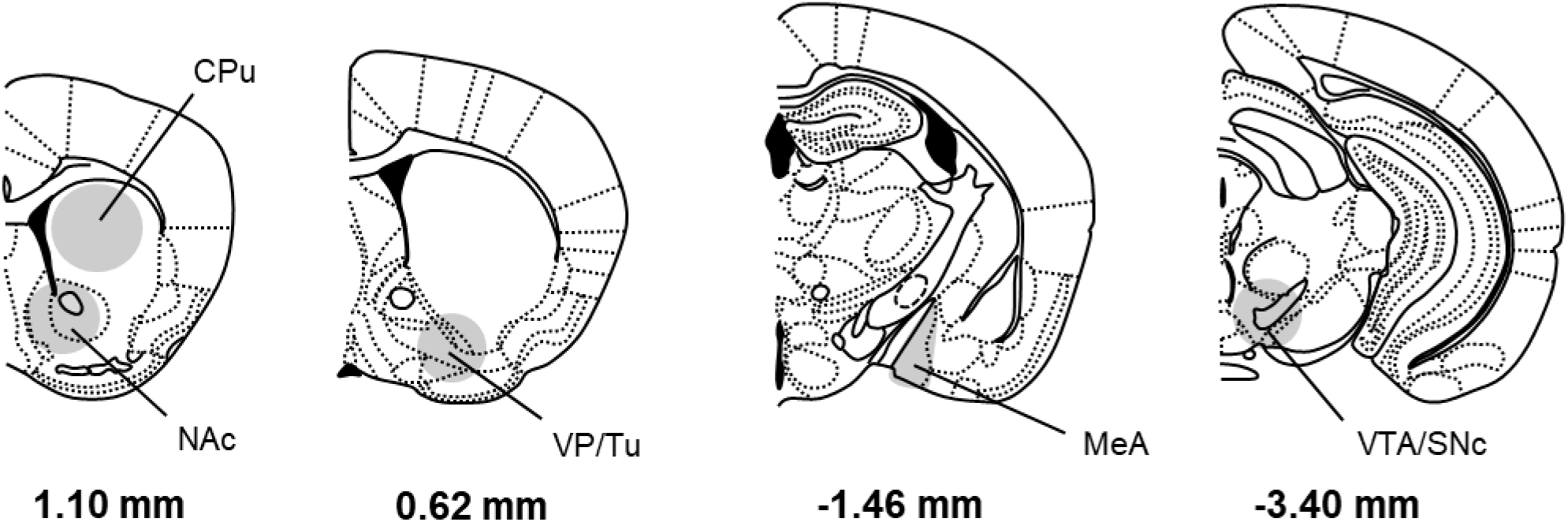
Schematic representations depict brain regions dissected for gene expression study and western blot experiments. CPu, NAc, VP/Tu and VTA/SNc were punched on 1-mm thick brain slices (CPu: one punch/side, ᴓ 2 mm; NAc, VP and VTA/SNc: one punch/side, ᴓ 1.25 mm). MeA was dissected out bilaterally. Coordinates refer to bregma. CPu: Caudate Putamen; NAc: Nucleus Accumbens; MeA: Medial Amygdala; SNc: Substancia Nigra, pars compacta; VP/Tu: ventral pallidum/olfactory tubercle; VTA/SNc: Ventral Tegmental Area/Substantia Nigra pars compacta.

**Figure S2.**
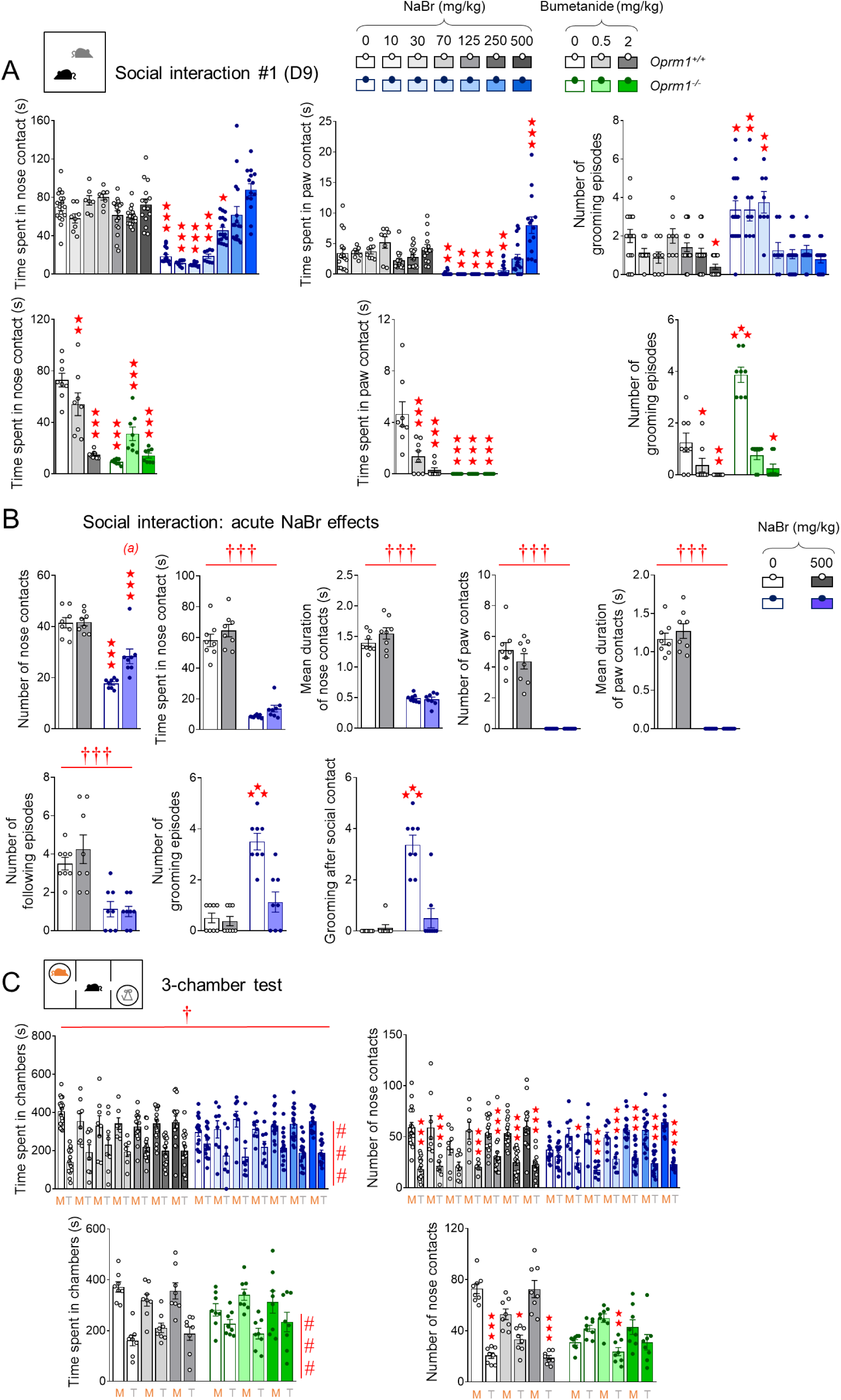
Chronic, and not acute, sodium bromide relieved social behavior deficits in *Oprm1^-/-^* mice, demonstrating superior effects to chronic bumetanide. See time line of experiments and animal numbers in Figure 1A, except for (B). (A) In the direct social interaction test (D9), chronic NaBr administration improved social interaction parameters in *Oprm1^-/-^* mice from the dose of 70 mg/kg but had no detectable effect in wild-type controls. Bumetanide effectively suppressed excessive grooming in these animals but demonstrated deleterious effects on social parameters in *Oprm1^+/+^* mice. (B) Acute administration of NaBr (250 mg/kg) in *Oprm1^-/-^* mice (versus *Oprm1^+/+^* mice, n=8 mice per genotype and treatment) partially increased the number of nose contacts and suppressed excessive grooming during direct social interaction but had no detectable effect on other parameters. (C) In the 3-chamber test, all mice spent globally more time in the chamber with the mouse versus the toy. NaBr treatment rescued social preference in *Oprm1* mutants since the dose of 10 mg/kg; bumetanide restored a preference for making more nose contacts with the mouse at the 0.5 mg/kg dose only. Results are shown as scatter plots and mean ± sem. Daggers: genotype effect, asterisks: treatment effect, solid stars: genotype x treatment interaction (comparison with wild-type vehicle condition), hashtag: stimulus effect (two-way ANOVA or three-way ANOVA with stimulus as repeated measure, followed by Newman-Keuls post-hoc test). One symbol: p<0.05, two symbols: p<0.01; three symbols: p<0.001.

**Figure S3.**
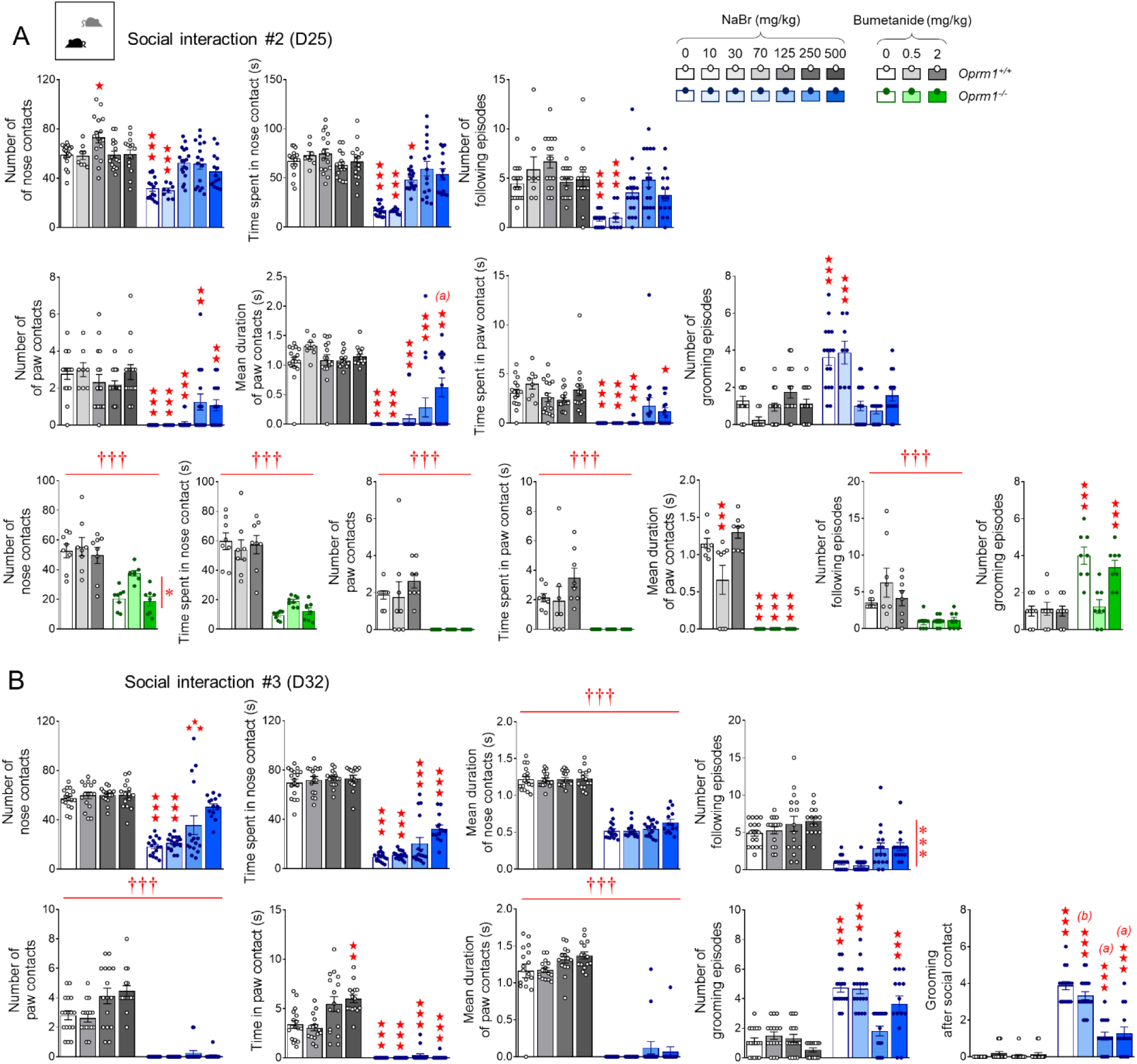
Chronic sodium bromide long-lastingly alleviated social interaction deficit in *Oprm1^-/-^* mice. See time line of experiments and animal numbers in Figure 1A. (A) One week after cessation of chronic treatment, beneficial effects of NaBr on social interaction parameters in *Oprm1^-/-^* mice were fully maintained for doses over 125 mg/kg. No significant effect was detected for bumetanide, except a reduction of grooming for the dose of 0.5 mg/kg. (B) Two weeks after cessation of NaBr treatment, an increase in the number and time spent in nose contacts was still detectable for the dose of 500 mg/kg in *Oprm1^-/-^* mice; grooming after social contact remained normalized from the dose of 250 mg/kg. Results are shown as scatter plots and mean ± sem. Daggers: genotype effect, asterisks: treatment effect, solid stars: genotype x treatment interaction (comparison with wild-type vehicle condition), (a) genotype x treatment interaction (comparison with knockout vehicle condition, p<0.001), (b) genotype x treatment interaction (comparison with knockout vehicle condition, p<0.01) (two-way ANOVA followed by Newman-Keuls post-hoc test). One symbol: p<0.05, two symbols: p<0.01; three symbols: p<0.001.

**Figure S4.**
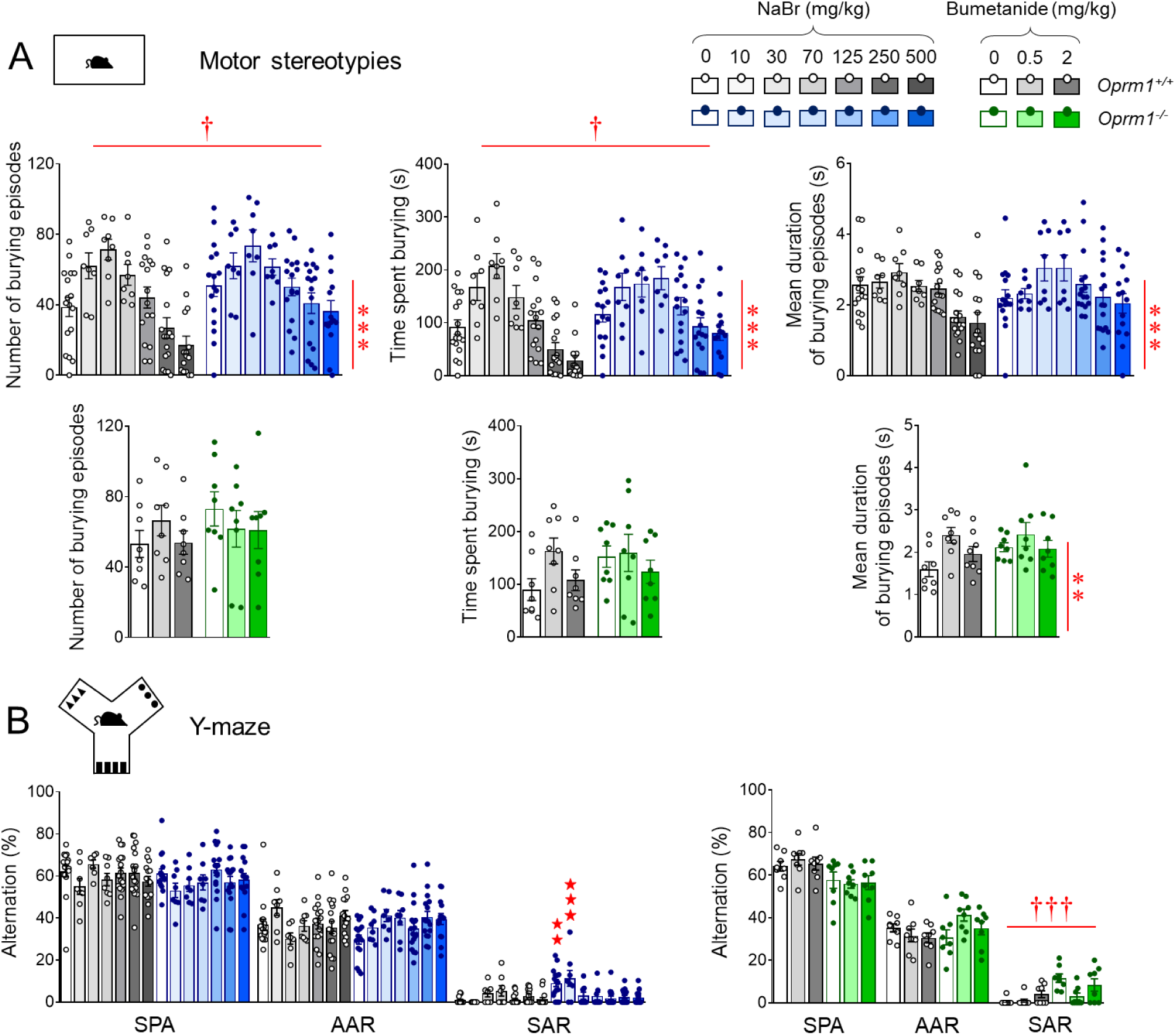
Chronic sodium bromide treatment reduced stereotypic behaviors and anxiety in *Oprm1^-/-^* mice. See time line of experiments and animal numbers in Figure 1A. (A) *Oprm1^-/-^* mice made more frequent and longer-lasting burying episodes than wild-type counterparts; bromide treatment increased these two behavioral outputs in both mouse lines. Chronic bromide and bumetanide increased the mean duration of burying episodes. (B) In the Y-maze, NaBr suppressed perseverative same arm returns in *Oprm1* null mice from the dose of 70 mg/kg without modifying spontaneous alternation or alternate arm returns; bumetanide had no significant effect in this test. Results are shown as scatter plots and mean ± sem. Daggers: genotype effect, asterisks: treatment effect, solid stars: genotype x treatment interaction (comparison with wild-type vehicle condition) (two-way ANOVA followed by Newman-Keuls post-hoc test). One symbol: p<0.05, two symbols: p<0.01; three symbols: p<0.001. AAR: alternate arm returns, SAR: same arm returns, SPA: spontaneous alternation.

**Figure S5.**
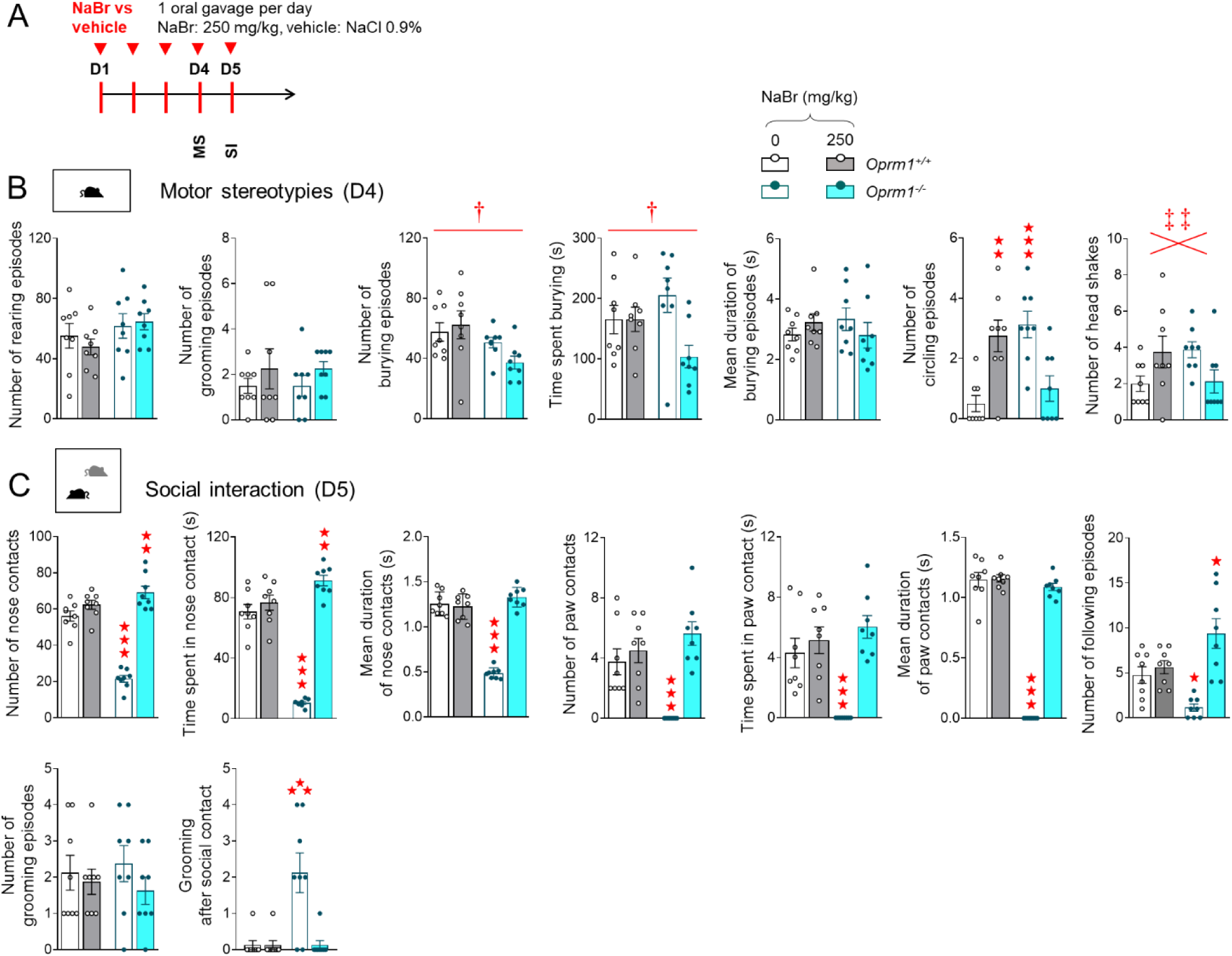
Subchronic oral administration of NaBr relieved autistic-like symptoms in *Oprm1^-/-^* mice. (A) *Oprm1^+/+^* and *Oprm1^-/-^* mice received either with NaBr (250 mg/kg) or vehicle (NaCl, 0.9%) via oral gavage once daily for 5 days (n=8 mice per genotype and treatment). Behavioral testing was performed on D4 and D5. (B) Oral subchronic NaBr administration suppressed stereotypic circling behavior in *Oprm1^-/-^* mice; it had opposite effects on the frequency of head shakes depending on mouse genotype, increasing this frequency in controls and decreasing it in mutant mice. (C) In the social interaction test, repeated oral NaBr treatment normalized behavioral parameters of *Oprm1* null mice to wild-type levels. Results are shown as scatter plots and mean ± sem. Daggers: genotype effect, double daggers: genotype x treatment interaction (post-hoc comparisons insignificant), solid stars: genotype x treatment interaction (comparison with wild-type vehicle condition) (two-way ANOVA followed by Newman-Keuls post-hoc test). One symbol: p<0.05, two symbols: p<0.01; three symbols: p<0.001. MS: motor stereotypies, SI: social interaction.

**Figure S6.**
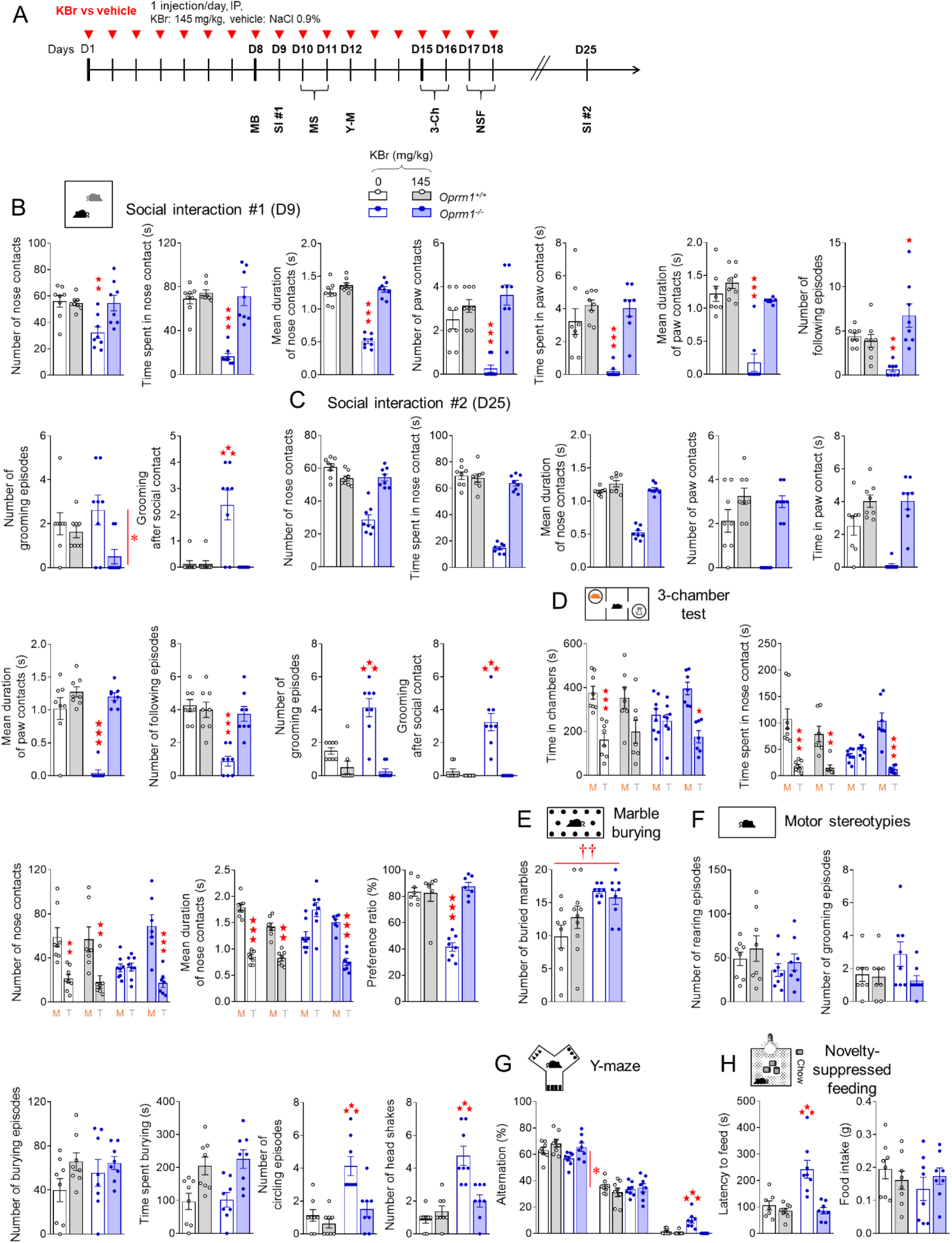
Chronic potassium bromide relieves autistic-like behaviors in *Oprm1^-/-^* mice. Chronic potassium bromide relieves autistic-like behaviors in *Oprm1^-/-^* mice. (A) *Oprm1^+/+^* and *Oprm1^-/-^* mice were treated either with KBr (0 or 145 mg/kg: n=8-10 mice per genotype and treatment) once daily for 18 days. Behavioral testing started on D8; social interaction was retested 1 week (D25) after cessation of chronic administration. (B) In the direct social interaction test (D9), chronic KBr relieved social deficits in *Oprm1* null mice without any detectable effect in *Oprm1^+/+^* controls. (C) This effect was maintained 1 week after cessation of chronic treatment. (D) In the 3- chamber test, KBr rescued preference for exploring a living congener over a toy in *Oprm1* mutants. As regards repetitive behaviors, KBr treatment (E) failed to reduce marble burying, (F) suppressed stereotypic circling and head shakes and (G) reduced perseverative same arm returns during Y-maze exploration in *Oprm1^-/-^* mice. (H) In the novelty-suppressed feeding test, potassium bromide normalized the latency to feed in *Oprm1* null mice without modifying food intake. Results are shown as scatter plots and mean ± sem. Daggers: genotype effect, asterisks: treatment effect, solid stars: genotype x treatment interaction (comparison with wild-type vehicle condition) (two-way ANOVA or three-way ANOVA with stimulus as repeated measure, followed by Newman-Keuls post-hoc test). One symbol: p<0.05, two symbols: p<0.01; three symbols: p<0.001. 3-Ch: 3-chamber test, AAR: alternate arm returns, M: mouse, MB: marble burying, MS: motor stereotypies, NSF: novelty-suppressed feeding, SAR: same arm returns, SPA: spontaneous alternation, SI: social interaction, T: toy, Y-M: Y-maze.

**Figure S7.**
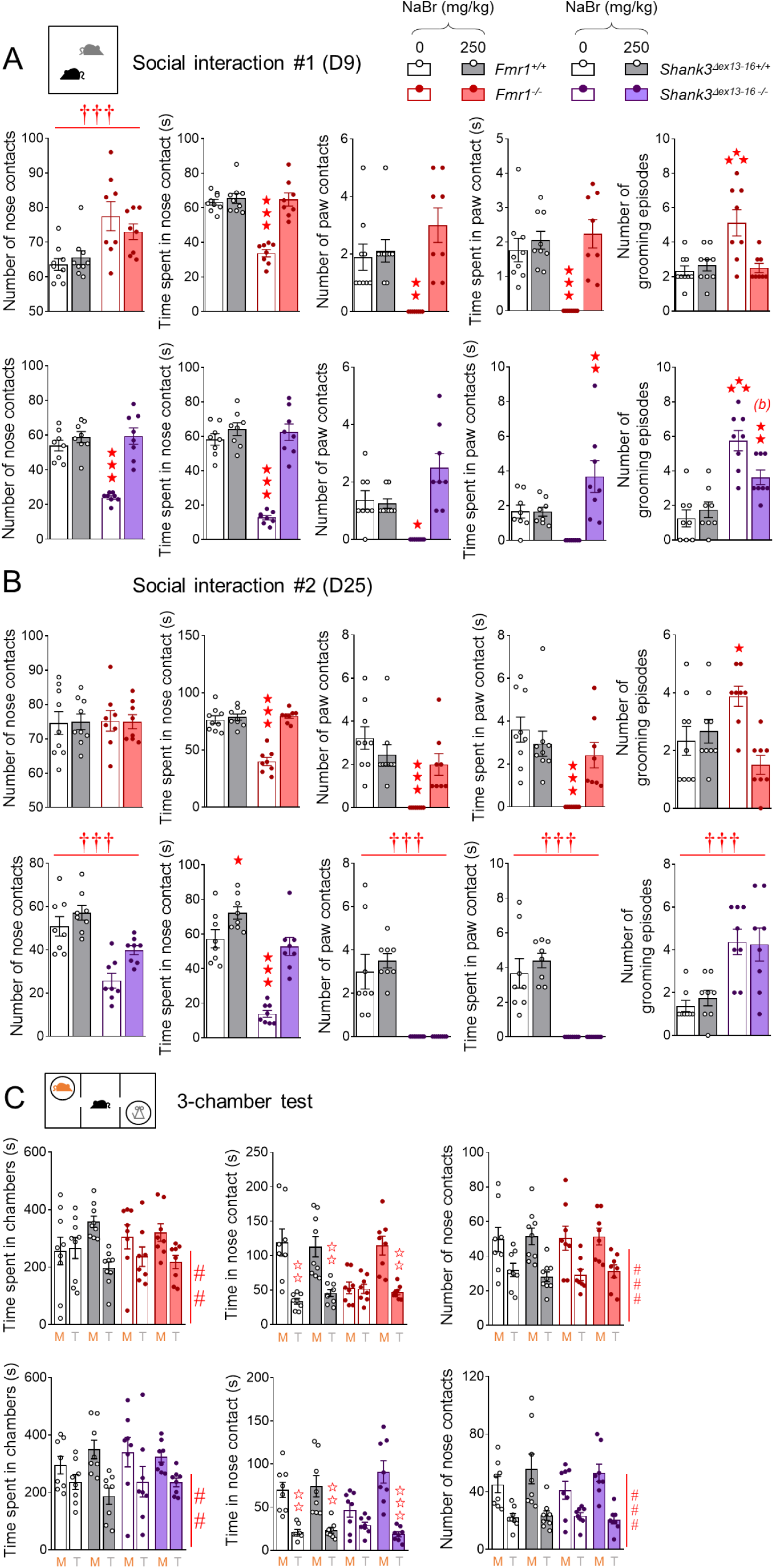
Figure 5. Chronic sodium bromide administration relieved social behavior deficits in *Fmr1^-/-^* and *Shank3^Δex13-16-/-^* mice. See animal numbers and time line of experiments in Figure 5. (A) Chronic NaBr treatment rescued direct social interaction (D9) in both *Fmr1* and *Shank3* mutant mouse lines. (B) Beneficial effects of NaBr treatment on social interaction parameters were fully maintained one week after cessation of treatment (D25) in in *Fmr1^-/-^* mice while only a restoration of the time spent in nose contact was detected in *Shank3^Δex13-16-/-^* mice. (C) In the 3-chamber test, all mice spent globally more time in the chamber with the mouse versus the toy. Results are shown as scatter plots and mean ± sem. Daggers: genotype effect, hashtag: stimulus effect, solid stars: genotype x treatment interaction (comparison to wild-type vehicle condition), open stars: genotype x treatment x stimulus interaction (mouse versus object comparison), (b) genotype x treatment interaction (comparison with knockout vehicle condition, p<0.01) (two-way ANOVA or three-way ANOVA with stimulus as repeated measure, followed by Newman-Keuls post-hoc test). One symbol: p<0.05, two symbols: p<0.01; three symbols: p<0.001. M: mouse, T: toy.

**Figure S8.**
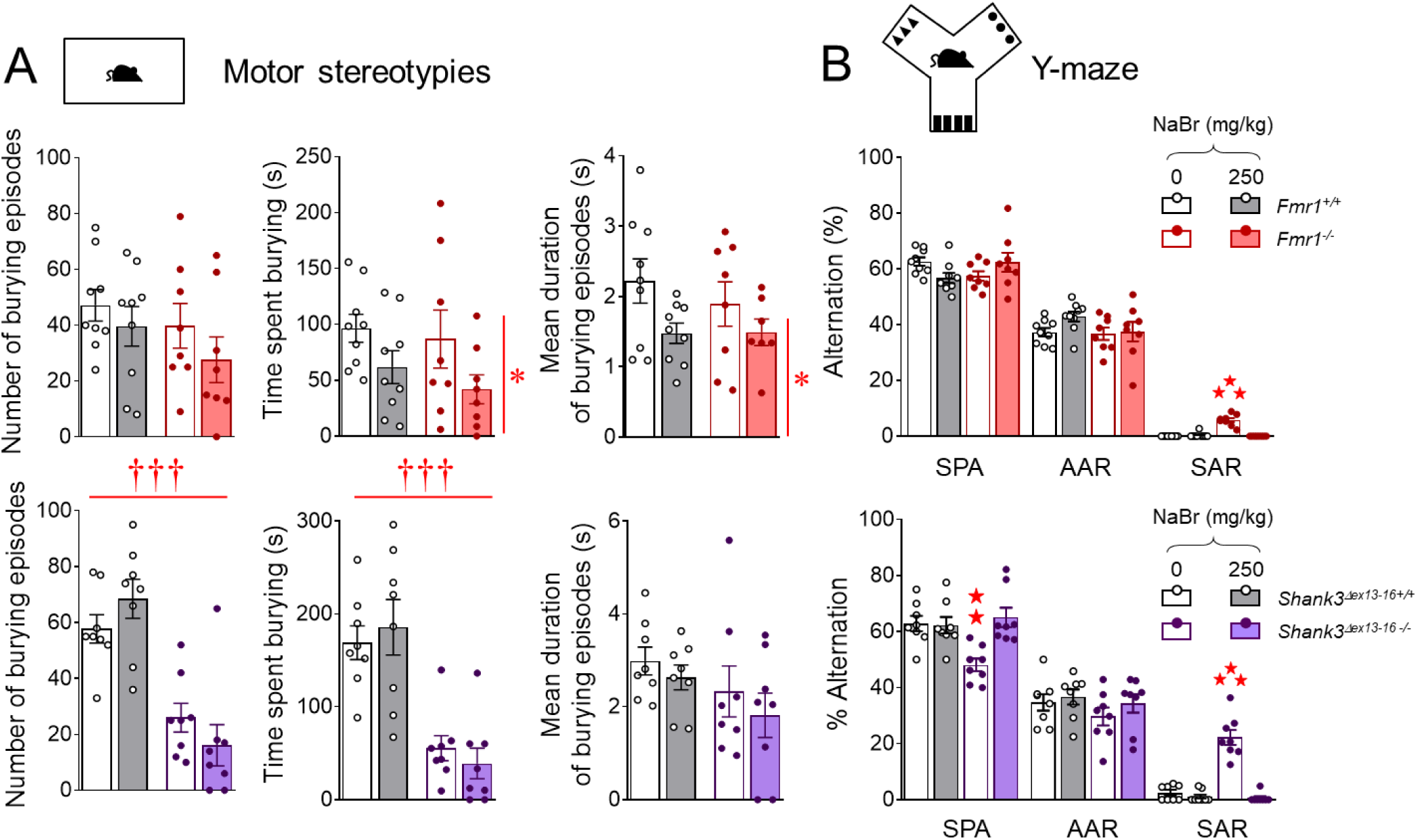
Chronic sodium bromide exposure relieved stereotypic behaviors and reduced anxiety levels in *Fmr1^-/-^* and *Shank3^Δex13-16-/-^* mice. See animal numbers and time line of experiments in Figure 5. (A) Chronic NaBr treatment decreased the time spent burying and mean duration of burying episodes in *Fmr1^-/-^* mice while it had no effect on deficient number of burying episodes and time spent burying in *Shank3^Δex13-16-/-^* mice. (B) In the Y-maze, chronic NaBr treatment normalized the percentage of preservative same arm returns in both mutant lines and restored spontaneous alteration rates in *Shank3^Δex13-16-/-^* mice. Results are shown as scatter plots and mean ± sem. Daggers: genotype effect, asterisk: treatment effect, solid stars: genotype x treatment interaction (comparison with wild-type vehicle condition) (two-way ANOVA followed by Newman-Keuls post-hoc test). One symbol: p<0.05, two symbols: p<0.01; three symbols: p<0.001.

**Figure S9.**
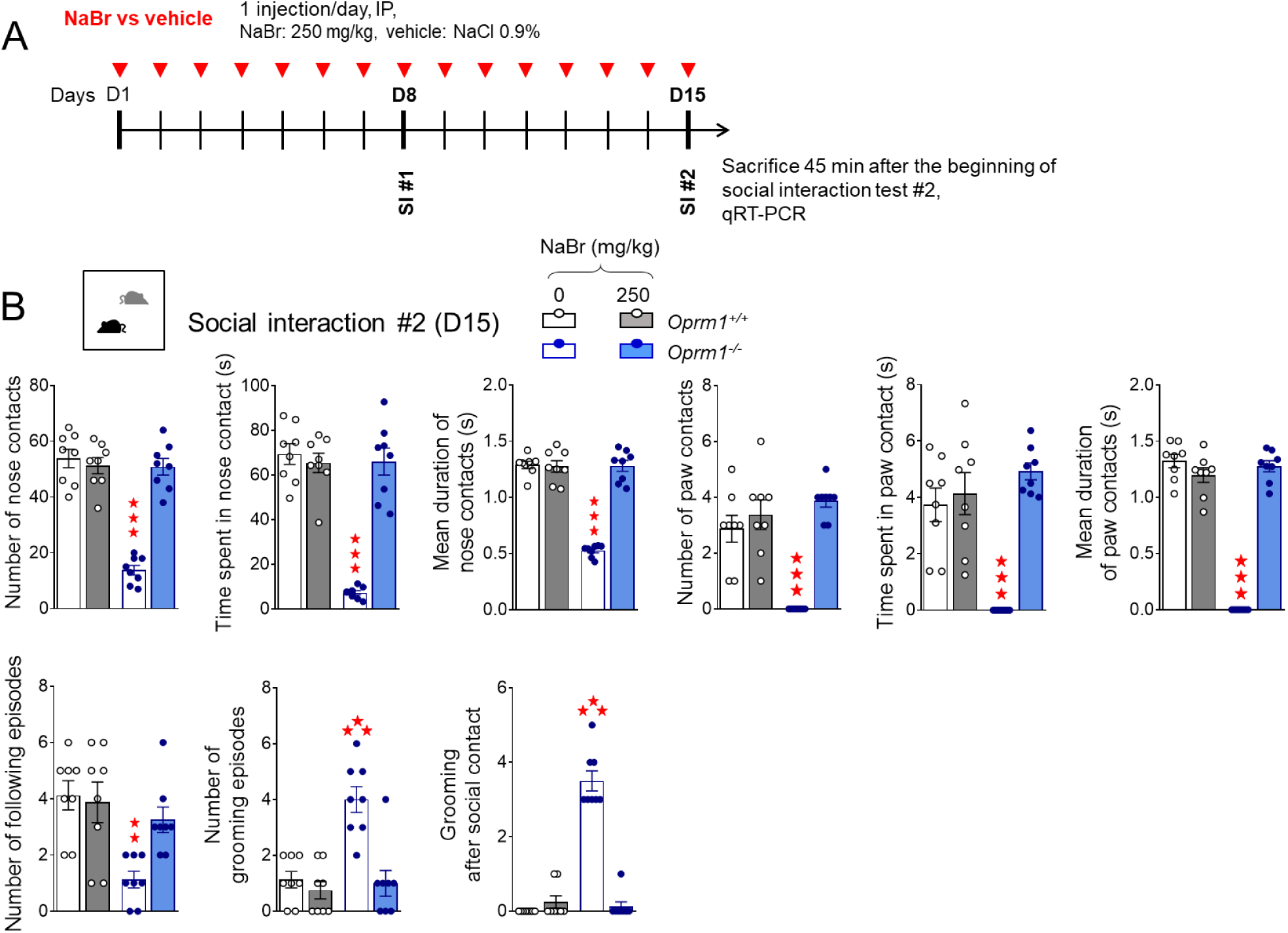
Effects of chronic NaBr treatment on social interaction and gene expression in *Oprm1^-/-^* mice. (A) *Oprm1^+/+^* and *Oprm1^-/-^* mice received either with vehicle or NaBr (250 mg/kg, i.p.) once daily for 15 days (n=8 mice per genotype and treatment). Mice were experienced social interaction on D8 and D15. Mice were sacrificed 45 min after the beginning of the second social interaction test. Behavioral parameters were assessed on D15, to allow correlations with gene expression data. (B) After two weeks of chronic treatment, NaBr at 250 mg/kg normalized social interaction parameters in *Oprm1^-/-^* mice to wild-type levels. Results are shown as scatter plots and mean ± sem. Solid stars: genotype x treatment interaction (comparison with wild-type vehicle condition) (two-way ANOVA followed by Newman-Keuls post-hoc test). Two symbols: p<0.01; three symbols: p<0.001.

**Figure S10.**
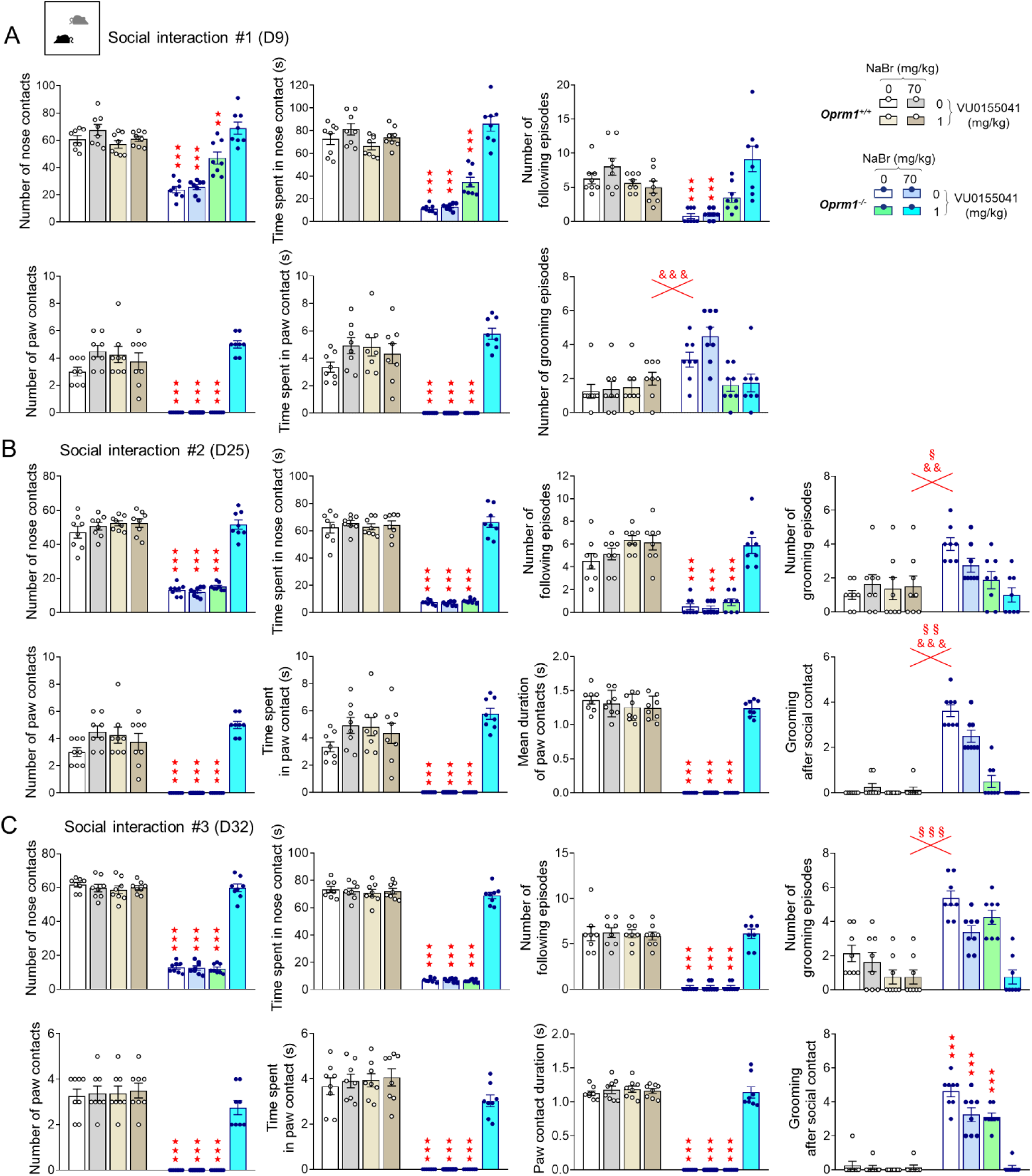
Long-lasting synergistic beneficial effects of sodium bromide and VU0155041 on social interaction in *Oprm1^-/-^* mice. See animal numbers and time line of experiments in Figure 8. (A) In the social interaction test (D9), synergistic beneficial effects of NaBr/VU0155041 combination treatment were observed in *Oprm1^-/-^* mice for all behavioral parameters, except for reducing grooming after social contact, for which VU0155041 (1 mg/kg) was sufficient to normalize behavior. These synergistic effects were fully maintained one week (B) and two weeks (C) after cessation of treatment. Results are shown as scatter plots and mean ± sem. Solid stars: genotype x NaBr x VU0155041 interaction (comparison to wild-type vehicle condition), ampersand: genotype x VU0155041 interaction, section: genotype x NaBr interaction (three-way or four-way ANOVA followed by Newman-Keuls post-hoc test). One symbol: p<0.05, two symbols: p<0.01; three symbols: p<0.001.

**Figure S11.**
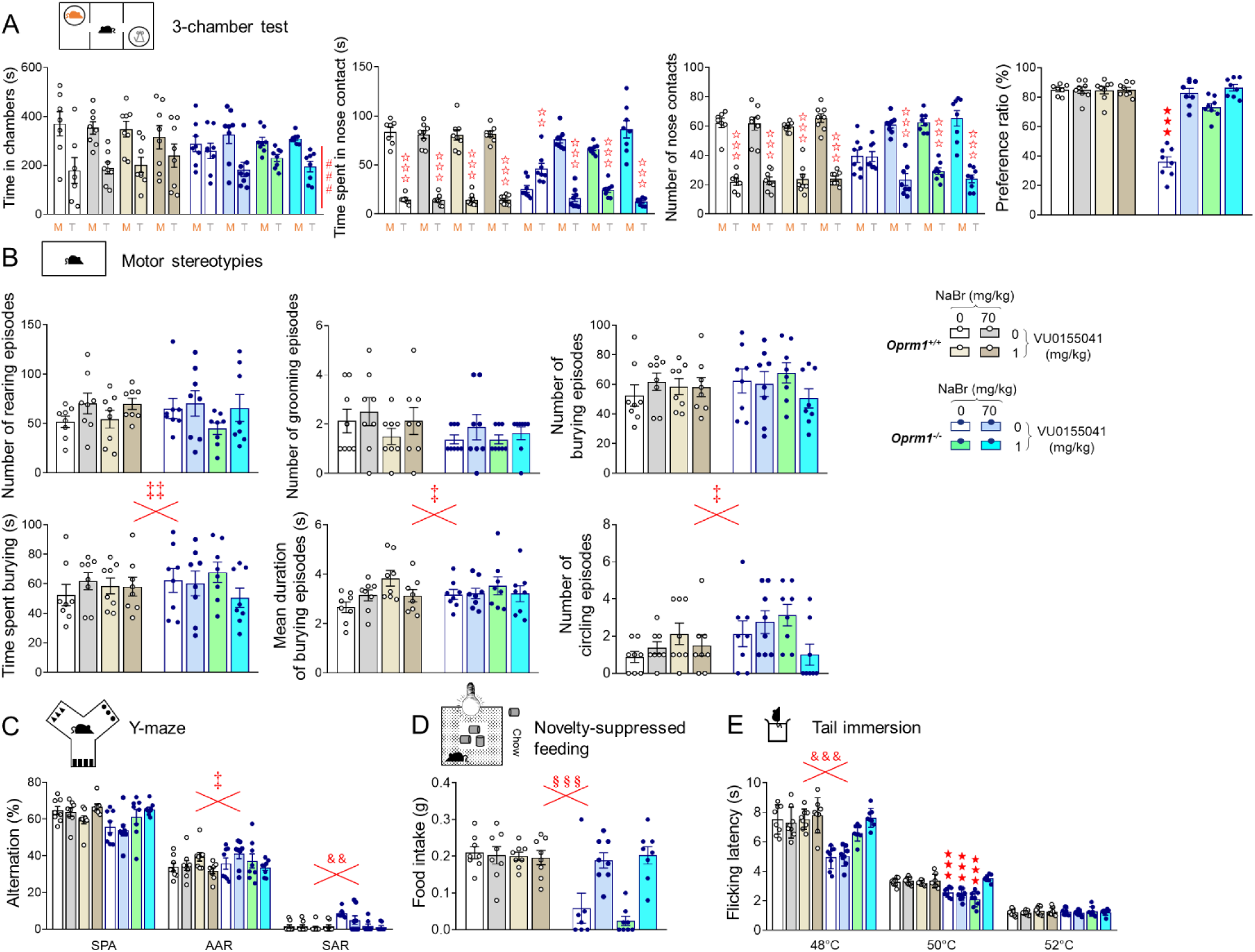
Synergistic beneficial effects of sodium bromide and VU0155041 on social preference and nonsocial behaviors in *Oprm1^-/-^* mice. See animal numbers and time line of experiments in Figure 8. (A) In the 3-chamber test, NaBr/VU0155041 combination had no effects on the time spent in chambers; NaBr (70 mg/kg) or VU0155041 (1 mg/kg) alone were able to fully restore other social preference parameters and preference ratio. (B) NaBr and VU0155041 treatments interacted to modulate time spent burying, duration of burying episodes and frequency of circling episodes independently from genotype. (C) In the Y-maze, NaBr and VU0155041 interacted to modulate the number of alternate arm returns while VU0155041 treatment was sufficient to suppress perseverative same arm returns in *Oprm1^-/-^* mice. (D) In the novelty-suppressed feeding test, NaBr administration was sufficient to restore food intake when the mice were back in their home cage. (E) In the tail immersion test, VU0155041 treatment on its own or in combination increased nociceptive thresholds in *Oprm1^-/-^* mice at 48°C while only combined NaBr/VU0155041 treatment restored these levels in mutants at 50°C. Results are shown as scatter plots and mean ± sem. Solid stars: genotype x NaBr x VU0155041 interaction (comparison to wild-type vehicle condition), open stars: genotype x stimulus x NaBr x VU0155041 interaction (mouse versus object comparison), double dagger: NaBr x VU0155041 interaction, ampersand: genotype x VU0155041 interaction, section: genotype x NaBr interaction (three-way or four-way ANOVA followed by Newman-Keuls post-hoc test). One symbol: p<0.05, two symbols: p<0.01; three symbols: p<0.001. AAR: alternate arm returns, M: mouse, SAR: same arm returns, SPA: spontaneous alternation, T: toy.

## Legends to Supplementary Tables

**Table S1.**
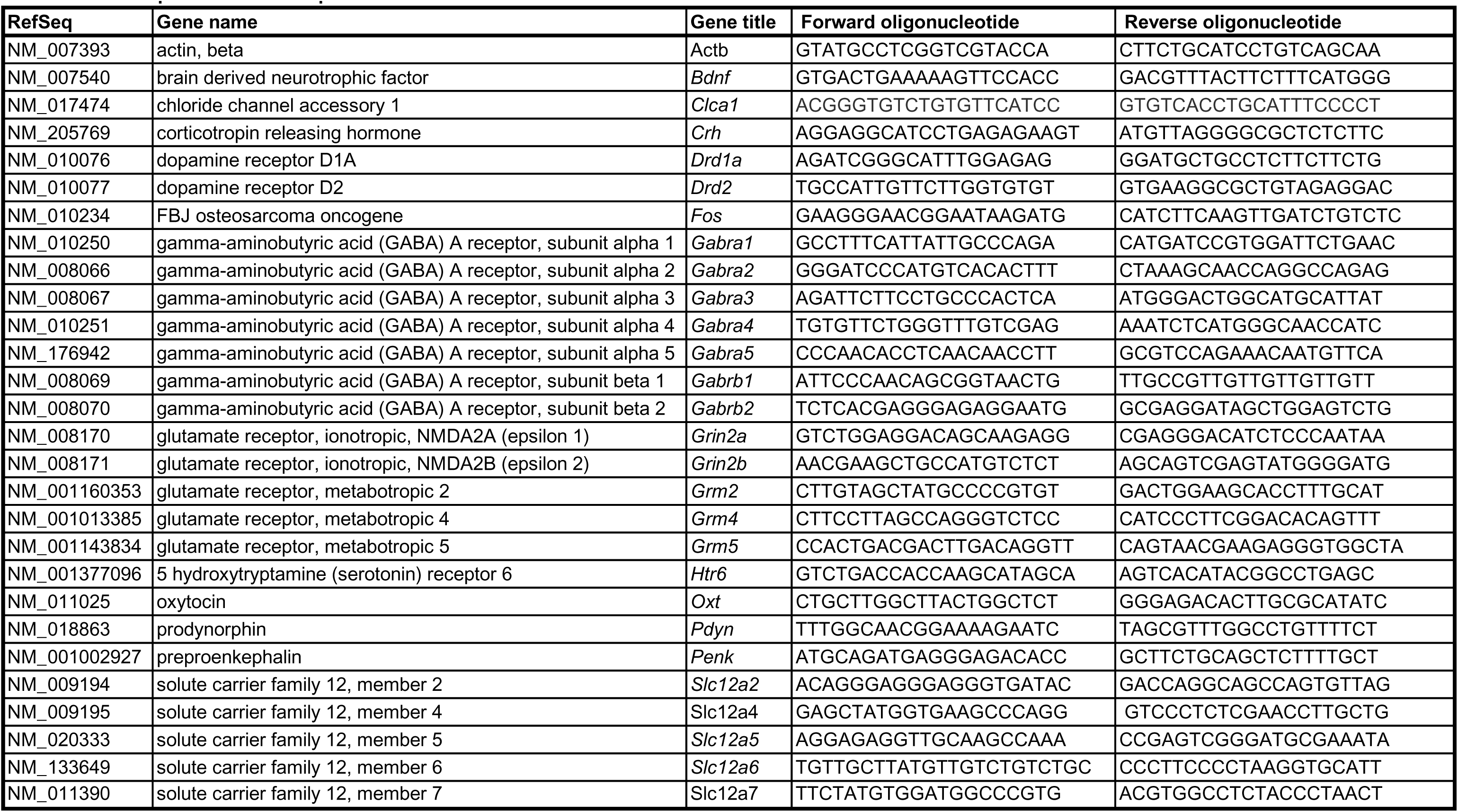
List of primers used for qRT-PCR.

**Table S2.**
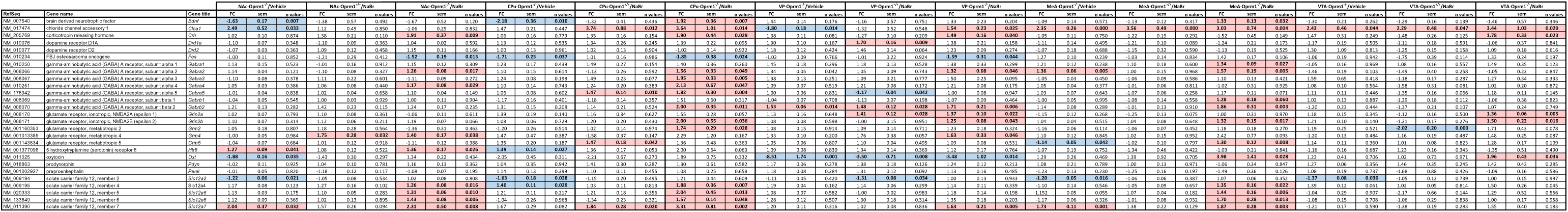
Transcription levels of a set of 27 genes in the CPu, NAc, VP/Tu, MeA and VTA/SNc in Oprm1^+/+^ or Oprm1^-/-^ mice treated chronically with NaBr versus vehicle. Data are expressed as fold change versus the vehicle - *Oprm1^+/+^*group (mean ± SEM). Two tailed t-tests were performed on transformed data (see Material and Methods). Significant regulations of gene expression are highlighted in bold and filled in red for significant up-regulation or in blue for significant down-regulation.

